# Dectin-1 Molecular Aggregation and Signaling is Sensitive to β-Glucan Structure and Glucan Exposure on *Candida albicans* Cell Walls

**DOI:** 10.1101/824995

**Authors:** Eduardo U. Anaya, Akram Etemadi Amin, Michael J. Wester, Michael E. Danielson, Kyle S. Michel, Aaron K. Neumann

## Abstract

Dectin-1A is a C-type Lectin innate immunoreceptor that recognizes β-(1,3;1,6)-glucan, a structural component of *Candida* species cell walls. The higher order structure of β-glucans ranges from random coil to insoluble fiber due to varying degrees of tertiary (helical) and quaternary structure. Model *Saccharomyces cerevisiae* β-glucans of medium and high molecular weight (MMW and HMW, respectively) are highly structured. In contrast, low MW glucan (LMW) is much less structured. Despite similar affinity for Dectin-1A, the ability of glucans to induce Dectin-1A mediated calcium influx and Syk phosphorylation positively correlates with their degree of higher order structure. Chemical denaturation and renaturation of MMW glucan showed that glucan structure determines agonistic potential, but not binding affinity, for Dectin-1A. We explored the role of glucan structure on Dectin-1A oligomerization, which is thought to be required for Dectin-1 signaling. Glucan signaling decreased Dectin-1A diffusion coefficient in inverse proportion to glucan structural content, which was consistent with Dectin-1A aggregation. Förster Resonance Energy Transfer (FRET) measurements revealed that molecular aggregation of Dectin-1 occurs in a manner dependent upon glucan higher order structure. Number and Brightness analysis specifically confirmed an increase in the Dectin-1A dimer and oligomer populations that is correlated with glucan structure content. Comparison of receptor modeling data with FRET measurements confirms that in resting cells, Dectin-1A is predominantly in a monomeric state. Super Resolution Microscopy revealed that glucan-stimulated Dectin-1 aggregates are very small (<15 nm) collections of a few engaged receptors. Finally, FRET measurements confirmed increased molecular aggregation of Dectin-1A at fungal particle contact sites in a manner that positively correlated with the degree of exposed glucan on the particle surface. These results indicate that Dectin-1A senses the solution conformation of β-glucans through their varying ability to drive receptor dimer/oligomer formation and activation of membrane proximal signaling events.

## Introduction

Overall, *Candida* infections have increased over the past 20 years in the United States [1–5]. It is estimated that 46,000 cases of healthcare-associated invasive candidiasis occur in the United States annually [6]. The fungal cell wall is composed of an inner layer of chitin, a middle layer of β(1,3;1,6)-D-glucan and an outer layer of N- and O-linked mannans [7]. During infection, the cell wall of *Candida* is an important and relevant virulence factor, playing roles in adhesion, colonization and immune recognition [8, 9].

Due to the abundant amount of mannan in the outer cell wall, β-glucan exhibits a very limited, punctate pattern of nanoscale surface exposure. The extent of this glucan masking is influenced by environmental conditions such as intestinal pH or lactate levels [10, 11]. In addition, interactions with neutrophils have been shown to “unmask” the mannose layer through a neutrophil extracellular trap-mediated mechanism [12]. Furthermore, our lab and others have determined that anti-fungal drugs “unmask” the fungal cell wall, which leads to increases in nanoscale regions of glucan exposure and correlates with enhanced host defense [13–15]. Therefore, fungal species use masking as a way to evade immune recognition of β-glucan by the host’s immune system [16].

β-glucans consist of a β-1,3-linked backbone with side chains of β-1,6-linked units that vary in length and degree of branching [17]. β-glucan forms triple-helical structures through intermolecular hydrogen bonds with two other strands [17–21]. This triple helix conformation is shown to form with just the β-1,3-linked backbone, however β-1,6-linked side chains play an important role in determination of the triple helix cavity formation through side chain/side chain interactions [21].

β-glucans are known for their biological activities such as enhancing anti-tumor, anti-bacterial, and anti-viral immunity as well as wound healing [22–25]. The biological activity of glucan is affected by its structure, size, structural modification, conformation and solubility [26]. Research has found that branching is not required to observe biological activity, but branching has been shown to enhance binding to the Dectin-1 receptor [27]. In contrast, β-glucan size is thought to play a major role in biological activity with glucans that are shorter than 10,000 Da being generally inactive *in vivo* [28, 29]. However, despite having similar sizes, glucans can display differences in their biological activities [30–32]. For example, studies have demonstrated that the immunoregulatory activity of variously sourced laminarin ranges from agonistic to antagonistic depending on its physicochemical properties, purity and structure [33]. Furthermore, β-glucans that have a triple helical conformation are more potent agonists of host immune response than single helical glucan [27,34–36]. We propose that the β-glucan triple-helix conformation plays an important role in determining the biological activity of the β-glucan through modulating the degree of receptor oligomerization upon ligation.

During innate immune recognition of *Candida*, the organization of cell wall ligands and pattern recognition receptors is an important determinant of successful immune activation [8]. The fungi-responsive C-type lectin (CTL) anti-fungal immunoreceptors play a central role in the detection of *Candida* [37]. Human Dectin-1A is the main CTL that recognizes the β-glucan found in the fungal cell wall [38–40]. Dectin-1A is found in myeloid lineage cells, and once activated, it stimulates phagocytic activity, the production of reactive oxygen intermediates and inflammatory mediators. Dectin-1A contains a CTL-like domain, separated from the cell membrane by a glycosylated stalk region, a transmembrane domain and an intracellular cytosolic domain. Dectin-1 contains half an Immunoreceptor Tyrosine-based Activation Motif (a YXXL sequence with an upstream stretch of acidic amino acids) in its cytoplasmic tail, which is termed a (hem)ITAM domain [41, 42]. Monophosphorylated ITAM domains, which are anticipated to approximate the structure of phosphorylated (hem)ITAM domains, poorly recruit and activate Syk for downstream signaling because they cannot support bivalent engagement of both of Syk’s SH2 domains [43]. Another (hem)ITAM bearing receptor, CLEC-2, is reported to require dimerization for its signaling [44]. By analogy to this and other (hem)ITAM receptors, it is speculated that Dectin-1A must oligomerize to recapitulate a multivalent binding site for Syk to facilitate signal transduction [8,44,45]. However, this prediction has not been directly explored for Dectin-1A in live cells at the molecular level with relation to signaling outcomes.

In this study, we propose that factors that induce an aggregated membrane organization of Dectin-1A during activation are very important for determining signaling outcomes of Dectin-1A engagement by β-glucan [44, 45]. We hypothesize that ligand structure, at the levels of glucan tertiary and quaternary structure as well as nanoscale glucan exposures on the pathogen surface, impacts signaling by determining the membrane organization and spatiotemporal clustering dynamics of Dectin-1A. To test this hypothesis, we stimulated HEK-293 cells transfected with Dectin-1A with a variety of soluble β-glucans that have different structures and sizes. We chose to work in this model system because it provides a simplified platform necessary to investigate the physical biology of Dectin-1A activation by isolating Dectin-1A signaling from the complex milieu of other receptors and other Dectin-1 isoforms expressed in innate immunocytes. Also, this model facilitates the expression of multiple fluorescent protein-tagged Dectin-1A constructs necessary to the work. Using calcium imaging and western blotting assays, our results revealed that Dectin-1A activation is influenced by the β-glucan triple helical structure. Furthermore, our subcellular FRET measurements by Fluorescence Lifetime Imaging Microscopy (FLIM-FRET), as well as application of fluorescence correlation spectroscopy approaches, revealed dimerization and oligomerization of Dectin-1A when stimulated with highly structured β-glucans. In addition, these dimerization events occurred in fungal contact sites of fungal cells with high glucan exposure. Together, our findings indicate β-glucan structure is required for Dectin-1A to form dimeric and oligomeric membrane aggregates.

## Results

### Dectin-1A activation is dependent on the molecular weight of the soluble β-glucan

β-glucans, existing as insoluble fibers in the cell wall, are likely to have a high degree of tertiary and quaternary structure. So, an encounter with highly structured glucan might be indicative of a pathogen cell wall structure. Furthermore, less structured glucans are encountered by Dectin-1A physiologically [41]. Small soluble circulating glucan can derive from sloughed cell wall material or from dietary absorption [46–49]. However, there is not much known about Dectin-1A’s ability to distinguish between highly structured β-glucan found on cell walls of fungal pathogens and less structured glucans found in circulation. Therefore, we examined how Dectin-1A activation is affected by glucans with different quaternary and tertiary structures. To accomplish this, we used high molecular weight (HMW 450 kDa), medium molecular weight (MMW 145 kDa), and low molecular weight (LMW 11 kDa) soluble glucans, in decreasing order of tertiary and quaternary structures, derived from *S. cerevisiae* cell walls. These *S. cerevisiae* glucans in soluble form or as particulate “zymosan” are common models for stimulation of innate immunocytes by fungal pathogen cell wall glucan. The above glucans have overall very similar composition and structures to *C. albicans* yeast glucan, though relatively minor differences in β-(1,6)-glucan side chain length and branching frequency have been reported between these species [50, 51].

Using these glucans, we performed intracellular calcium ([Ca^2+^]_i_) measurement experiments on HEK-293 cells (lacking endogenous Dectin-1 expression) transfected with Dectin-1A. We stimulated the cells using either LMW, MMW, or HMW glucan. All the glucans induced a significant increase in peak amplitude [Ca^2+^]_i_ compared to unstimulated cells (Fig.1 A,B). We found that large, highly ordered glucans (HMW and MMW) induced a significant, Dectin-1 dependent increase in peak amplitude of [Ca^2+^]_i_ compared to LMW, with MMW having the highest peak amplitude (Fig. 1C). Interestingly, the response to MMW glucan was uniform at the single cell level. However, cells stimulated with HMW glucan exhibited a more heterogeneous response, with some cells achieving comparable maximum amplitudes as with MMW and others exhibiting little change from basal calcium levels (Fig. 1A, B). We expect that HMW glucan is present as larger particles than MMW, so at equal mass/volume concentrations, the HMW solution will have a lower concentration of particles. Non-responder cells in HMW experiments may have stochastically encountered too few glucan particles to achieve a detectable signaling response. When we stimulated cells with MMW or HMW at equimolar concentrations, which should have similar glucan particulate concentrations, we observed a minor difference in peak amplitude, but we saw that the integrated Ca^2+^ flux over time was the same for MMW and HMW glucans (Supplemental Fig. 1). Furthermore, single cell data demonstrated a similarly uniform response of Dectin-1 to MMW and HMW glucans under these conditions. These results indicate that Dectin-1A drives differential Ca^2+^ flux to glucan ligands that vary in size and structure.

**Figure 1.**
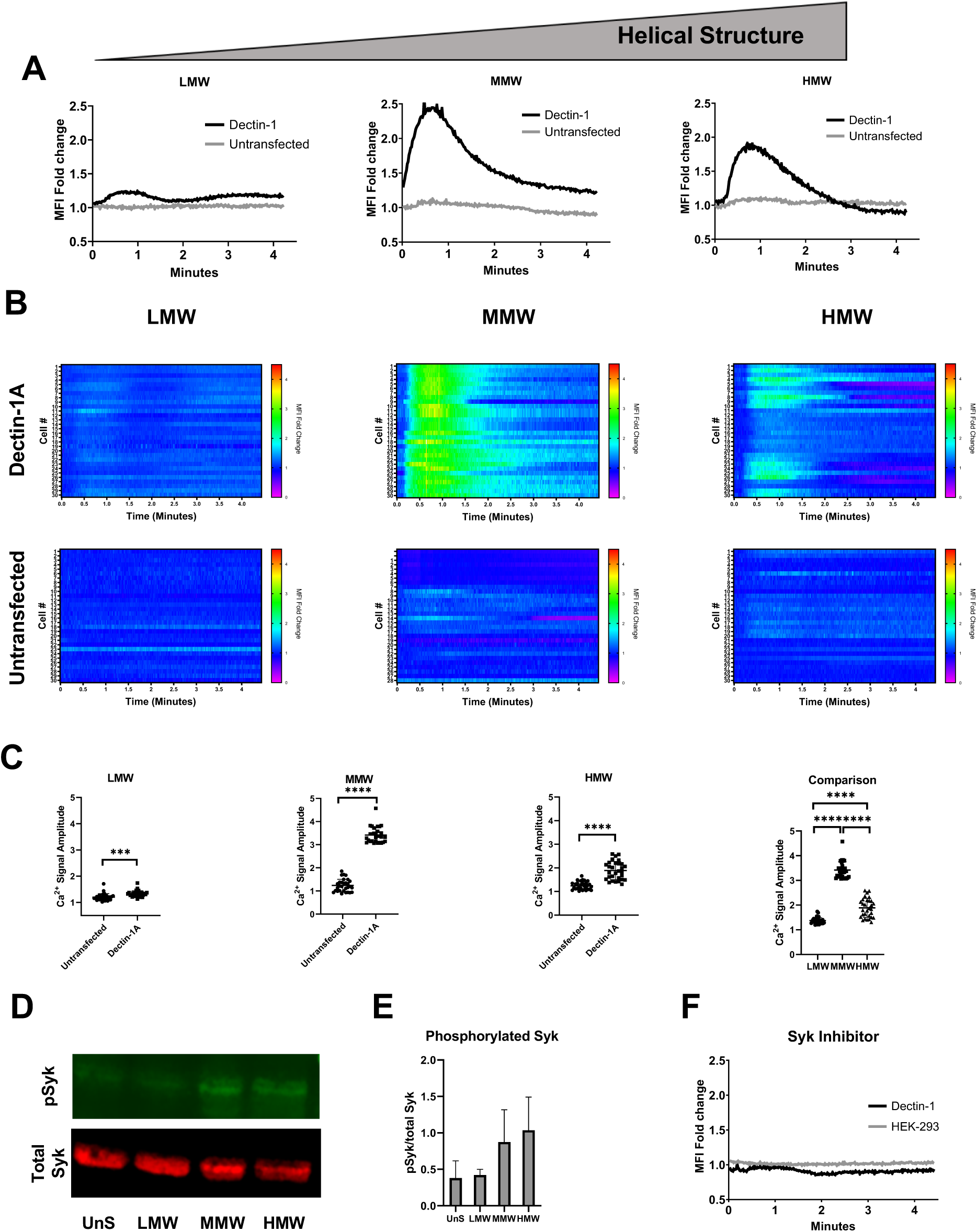
Differential signaling response of Dectin-1A to soluble β-glucan. (A) HEK-293 cells stably transfected with Dectin-1A were loaded with Fluo-4 and Cell Tracker Deep Red at equimolar concentrations. Cell Tracker Deep Red was simultaneously loaded in order to normalize for changes in cytosolic volume caused from cell contraction. The mean fluorescence intensity of 30 cells was averaged for Dectin-1A transfected (black) or untransfected (grey) HEK-293 cells stimulated with LMW, MMW, or HMW glucan at 1 µg/ml. (n = 30 per glucan from 3 independent experiments per glucan). Data shown as mean fold change in volume-normalized [Ca^2+^]_intracellular_. (B) Heat maps of relative changes in intracellular calcium concentration of untransfected or Dectin-1A transfected individual cells upon addition of either LMW, MMW, or HMW glucan. Each row represents the normalized ratio of Fluo-4 and Cell Tracker Deep Red for a single cell over time. (C) Maximum signal amplitude of single cells treated with LMW, MMW, or HMW glucan showing Dectin-1 dependent glucan responses. Statistical comparison of maximum amplitudes for Dectin-1 expressing cells treated with each of the three glucans is shown in the far right panel. Data shown as mean ± SD (n = 30 per glucan from 3 independent experiments per glucan). One-way ANOVA with multiple comparisons by the Dunnett test, *** p<0.0001, **** p<0.000001. (D) Cell lysates were collected at 5 min after stimulation and analyzed by Western blotting using antibodies against p-Syk and Syk. The intensity of p-Syk was normalized against total Syk. (E) Quantification of the western blot normalized p-Syk/Syk signal is shown as mean ± SD of n = 3 independent experimental replicates. (F) Untransfected (grey) and stably transfected Dectin-1A (black) HEK-293 cells were loaded with Fluo-4 and Cell Tracker Deep Red at equimolar concentrations and treated with Syk Inhibitor at 250 nM, then stimulated with MMW glucan. Average mean fluorescence intensity of 30 cells was observed for cells stimulated with MMW glucan at 1 µg/ml. (n = 30 from 3 independent experiments).

To determine how these differently-structured, soluble glucans impacted cellular patterns of Syk phosphorylation, we stimulated HEK-293 cells expressing Dectin1A-mEmerald with H_2_O (vehicle), LMW, MMW, or HMW. Whole cell lysate was collected and Syk phosphorylation was determined by western blot analysis. Likewise, our results show an increase in Syk phosphorylation in the larger, highly structured glucan MMW and HMW compared to unstimulated and LMW stimulated cells (Fig. 1 D,E).

Additionally, Syk inhibitors abrogated calcium signaling of Dectin-1A when stimulated with MMW glucan (Fig. 1F). These results indicate that glucans with higher order structure are better able to activate Dectin-1A-mediated Ca^2+^ signaling and that this is a Syk dependent process.

### β-glucan denaturation abrogates its potential for Dectin-1A activation

To determine if the glucan structure affects signaling outcomes, we denatured MMW (highly stimulatory glucan) using DMSO, a chaotrope that promotes a reduction in glucan tertiary structure, thus shifting MMW’s triple helix structure to a more single helical or random coil structure [17]. The results showed that when we denatured MMW glucan, we did not observe calcium signaling in cells expressing Dectin-1A (Fig. 2A, B). However, when we renatured the glucans by removing DMSO via dialysis we observe partial recovery of calcium signaling. We found that renatured glucans induce a significant increase in peak amplitude [Ca^2+^]_i_ response compared to DMSO denatured MMW and renatured MMW stimulated untransfected HEK-293 cells (Fig. 2 C). In addition, we confirmed the loss of helical structure via a Congo Red assay. Congo Red specifically binds to β-(1,3)-glucans with a triple helix conformation as their tertiary structure. This binding is detected by bathochromic shift in absorbance maximum from 488 to 516 nm [52]. Our results indicated a loss in glucan structure after denaturation in a DMSO solution that was regained upon renaturation (Fig. 2 D). Moreover, we repeated these experiments by stimulating cells with glucan denatured with NaOH or neutralized renatured glucan [17]. Similarly, our results show that cells lose the ability to activate Dectin-1A calcium signaling when stimulated with denatured MMW glucan but regain the ability to stimulate Dectin-1A activation when the glucan is renatured (Fig. 2 E,F). We found that renatured glucans induce a significant increase in peak amplitude [Ca^2+^]_i_ response compared to NaOH denatured MMW and renatured MMW stimulated untransfected HEK-293 cells (Fig. 2 G). In addition, we confirmed that glucan structure was lost when NaOH was added and regained when neutralized (Fig. 2 H). These results suggest that glucan structure is an important factor in activating a Dectin-1A response.

**Figure 2:**
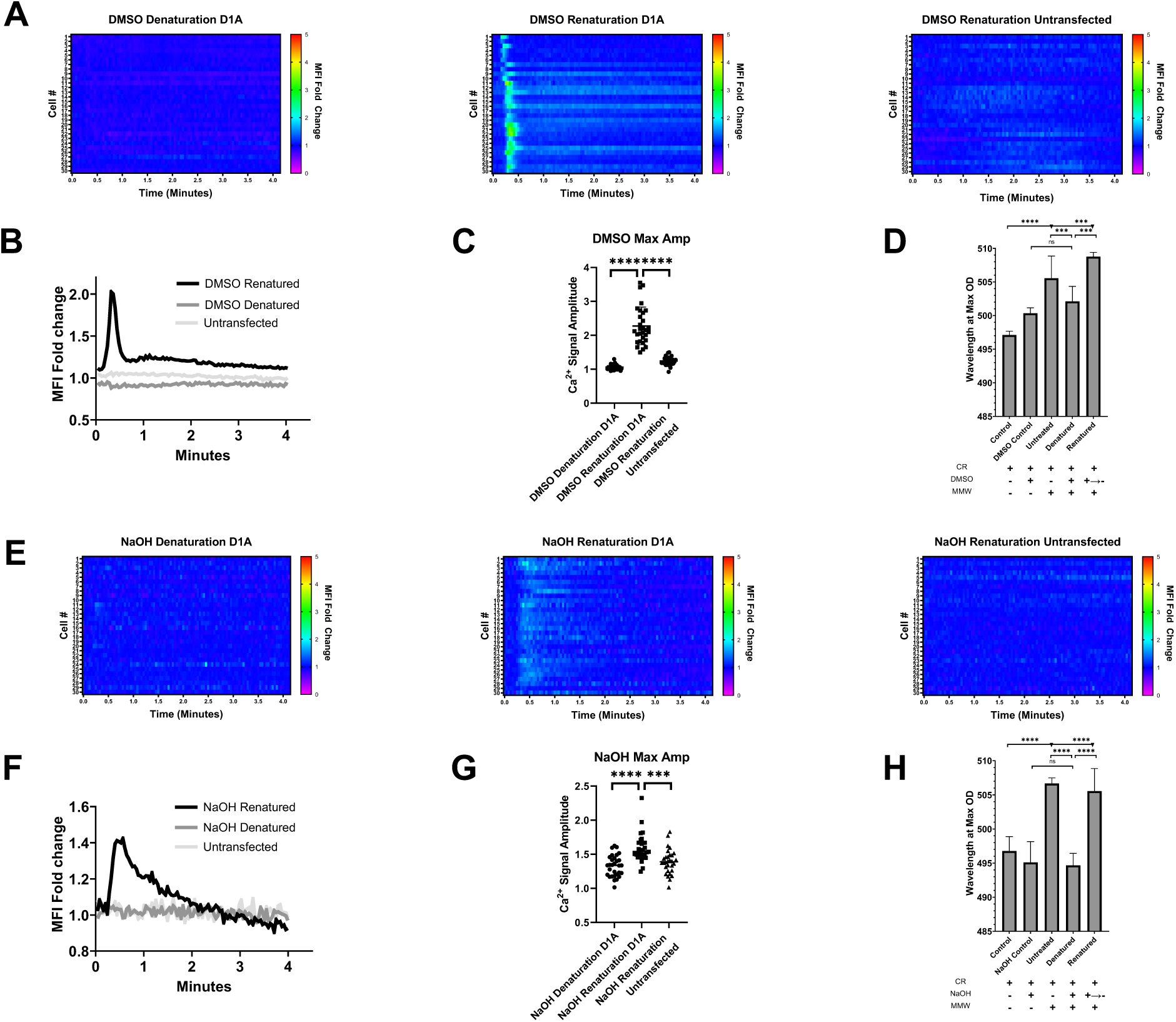
Glucan higher order structure is a critical determinant of its stimulatory potential. (A) HEK-293 cells stably expressing Dectin-1A or untransfected were loaded with Fluo-4 and Cell Tracker Deep Red at equimolar concentrations. Heat maps show relative changes in intracellular calcium concentration of untransfected or Dectin-1A transfected individual cells upon either addition of DMSO denatured MMW glucan or renatured MMW glucan. Each row represents the normalized ratio of Fluo-4 and Cell Tracker Deep Red for a single cell over time. (B) Average mean fluorescence intensity of 30 cells stimulated with DMSO denatured/renatured MMW glucan was observed. (n = 30 from 3 independent experiments). (C) Maximum amplitude of single cells treated with DMSO denatured or renatured MMW glucan. Data shown as mean ± SD (n = 30 from 3 independent experiments). One-way ANOVA with multiple comparisons by the Dunnett test, **** p<0.000001. (D) 1 mg/ml of MMW glucan was denatured using DMSO and incubated with Congo Red. Control samples contained Congo Red and DMSO. Renaturation was accomplished by dialyzing out DMSO 24 hrs prior to the experiment. Data shown as mean ± SD (n = 9 from 3 independent experiments). (E) HEK-293 cells stably expressing Dectin-1A or untransfected were loaded with Fluo-4 and Cell Tracker Deep Red at equimolar concentrations. Heat maps show relative changes in intracellular calcium concentration of untransfected or Dectin-1A transfected cells upon either addition of NaOH denatured or renatured MMW glucan. Each row represents the normalized ratio of Fluo-4 and Cell Tracker Deep Red for a single cell over time. (F) Average mean fluorescence intensity of 30 cells stimulated with NaOH denatured/renatured MMW glucan was observed. (n = 30 from 3 independent experiments). (G) Maximum amplitude of single cells treated with NaOH denatured or renatured MMW glucan. Data shown as mean ± SD (n = 30 from 3 independent experiments). One-way ANOVA with multiple comparisons by the Dunnett test, **** p<0.000001. (H) 1 mg/ml of MMW glucan was denatured using NaOH and incubated with Congo Red. Control samples contained NaOH and Congo Red. Renaturation was accomplished by neutralizing NaOH. Data shown as mean ± SD (n = 9 from 3 independent experiments). Welch’s t-test, ** p<0.001, *** p<0.0001, **** p<0.000001.

### β-glucan structure variation and manipulation does not alter affinity for Dectin-1

We considered the possibility that these soluble glucans might have different affinities for Dectin-1A, resulting in differential receptor activation. Thus, we conducted biolayer interferometry experiments to determine the binding affinity of these glucans to the carbohydrate recognition domain of Dectin-1A. This was accomplished using a chimeric fusion protein of the carbohydrate recognition domain of Dectin-1A and the human IgG Fc region. An anti-human IgG Fc Capture biosensor tip was used to load this fusion protein. Association and dissociation curves of the glucan and fusion protein where then collected. The results shown in Fig. 3A indicate that all the glucans have approximately nanomolar dissociation constants for Dectin-1A carbohydrate recognition domain despite the previously described differences in signaling [27]. Weight average molecular weights of purified *Saccharomyces cerevisiae* β -(1,3)-glucan fractions vary over an approximate 40-fold range, but there is relatively little difference between these glucans’ apparent affinity for the Dectin-1 carbohydrate recognition domain.

**Figure 3:**
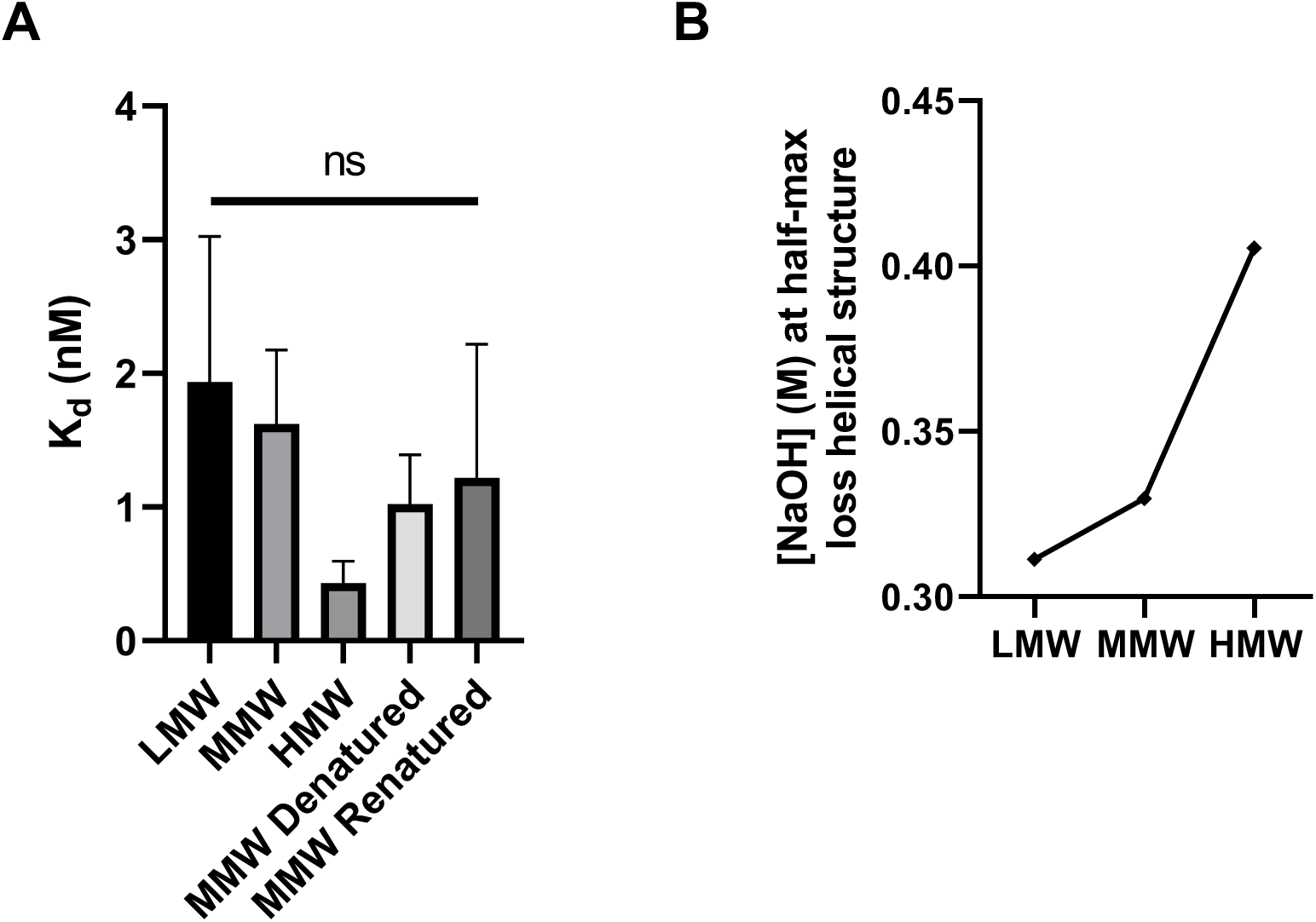
Characterization of Dectin-1 binding affinity and helical structure of fungal cell wall glucans used in this study. (A) Biolayer interferometry experiments were conducted on LMW, MMW, HMW, MMW denatured, and MMW renatured glucans using an anti-human IgG Fc Capture biosensor tip and a Dectin-1A-Fc fusion protein. Data shown as mean ± SD (n = 3 from 3 independent experiments). Statistical comparison by one-way ANOVA. (B) LMW, MMW, and HMW β-glucans were denatured using 0M-1M NaOH in the presence of Congo Red. Concentration of NaOH at which absorbance (516 nm) decreased to the half-maximal value was plotted. Data shown as mean ± SD (n = 9 from 3 independent experiments).

Furthermore, to determine differences in the structure of these glucans, we analyzed the conformational transition of triple helix to random coil of β-1,3-D-glucans through denaturation experiments. Experiments were conducted by denaturing glucans with NaOH at various concentrations in the presence of Congo Red. Our results show that the amount of glucan tertiary structure scales with molecular weight as measured by the concentration of NaOH required to reduce Congo Red binding to glucan (Fig. 3B), suggesting that the size of the glucans is correlated with their higher-order structure. Together, these results indicate that downstream signaling of the receptor is determined by the structure of the glucan rather than affinity alone.

### Dectin-1A decreases in diffusion coefficient when stimulated with highly structured β-glucans

The Stokes-Einstein equation predicts that if a diffusing object increases in hydrodynamic radius, it will slow down proportionally to that change. We measured the diffusion coefficient of Dectin-1A pre/post glucan stimulation to determine whether the receptor diffusion coefficient decreased, potentially due to increasing hydrodynamic radius as an initially monomeric receptor formed larger clusters/oligomers. We obtained average diffusion coefficients and spatial number density of our receptor using Raster Image Correlation Spectroscopy (RICS). RICS allowed us to survey multiple areas of the cell for molecular parameters such as diffusion coefficient and receptor density. This section pertains to receptor diffusion measurement by RICS, while receptor density is treated in a separate section below. Furthermore, because fluorescence was probed within a large cell area, RICS analyses suffered much less from photobleaching and location specific artifacts than analogous single-point measurements. Previous research has described RICS in more detail [53, 54]. Briefly, we generated a volume of excitation using focused laser illumination and calibrated a confocal observation volume using standard fluorophores of known diffusion coefficient. We observed fluorescent molecules (i.e., Dectin-1A-mEmerald) diffusing in and out of this excitation volume through fluctuation in the number of photons obtained. Experimentally observed fluorescence correlations at various spatiotemporal lags were then fit to a 2D autocorrelation function to obtain the receptor diffusion coefficient in the observed membrane (Fig. 4A).

**Figure 4:**
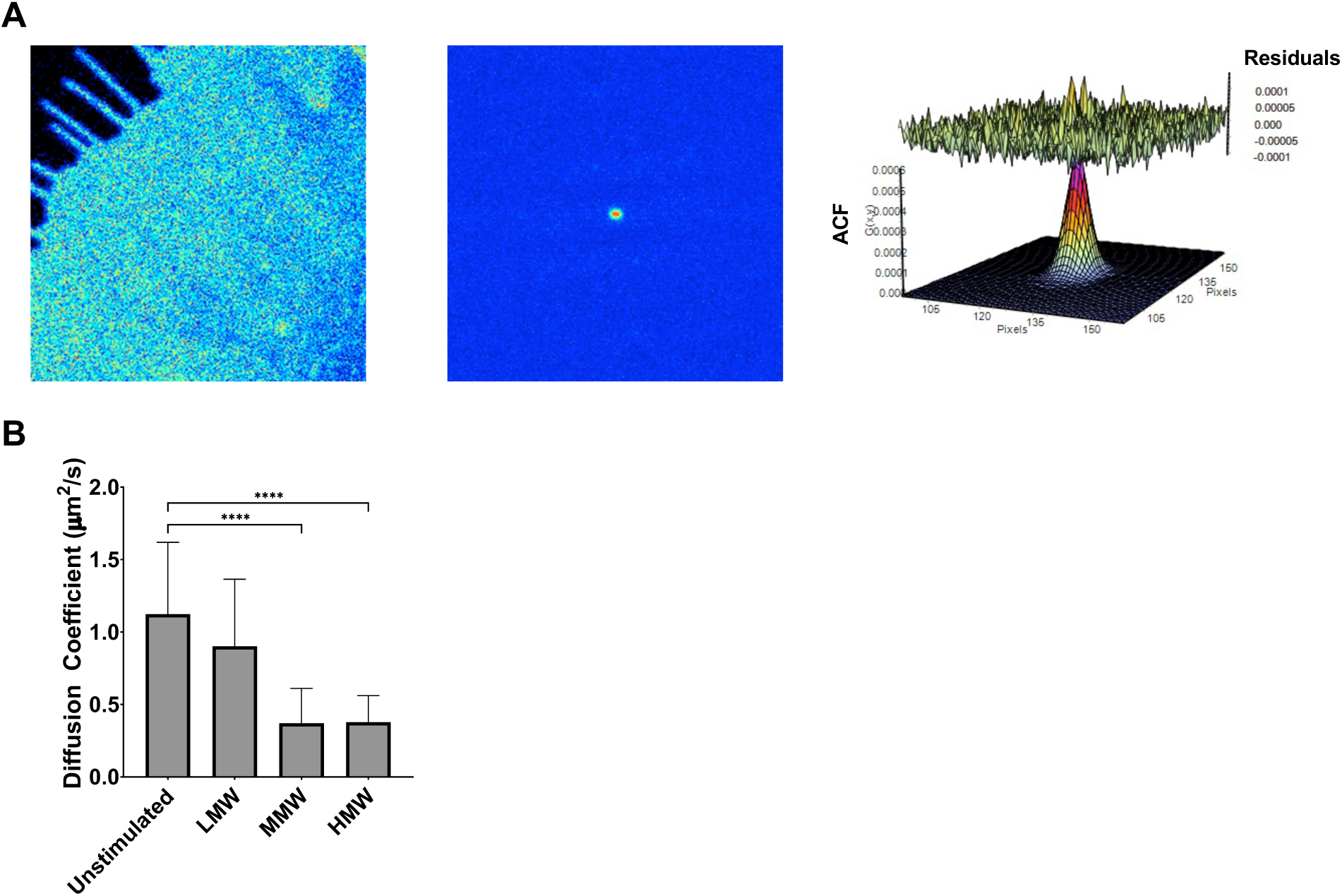
Dectin-1A surface diffusion coefficient decreases when stimulated with highly structured β-glucans. (A) A representative example of Raster Image Correlation Spectroscopy (RICS) analysis. (Left) Representative RICS image of HEK-293 cells expressing Dectin-1A-mEmerald. (Middle) Autocorrelation function (ACF) calculated from the time series. Red represents a high ACF value, blue represents a low ACF value. (Right) Fit of the ACF to a Gaussian diffusion model to calculate the diffusion coefficient. (B) RICS analysis of fluorescently tagged Dectin-1A expressed in HEK-293 provided average diffusion coefficient for cells that were unstimulated or stimulated with LMW, MMW, or HMW. Data shown as mean ± SD (n = 30); One-Way ANOVA multiple comparison test, **** p<0.00001.

Using HEK-293 cells expressing Dectin-1A-mEmerald, we conducted RICS measurements before and after stimulation with soluble β-glucans. We determined that cells stimulated with MMW or HMW exhibited a significant decrease in mobility compared to LMW and unstimulated cells (Fig. 4B). This finding is consistent with an increase in receptor aggregation upon stimulation, which we examine in greater detail below. We proceeded to conduct additional membrane biophysical studies to further test for changes in the molecular aggregation state of Dectin-1 during stimulation with glucan.

### Dectin-1A forms dimers/oligomers when stimulated with highly structured β-glucans

The results shown above indicate that the β-glucan structure is an important factor in signaling outcomes. Previous research has shown that other transmembrane CTLs that also contain a (hem)ITAM domain can form homodimers before or upon ligand recognition [55, 56]. Furthermore, crystallography studies of the carbohydrate recognition domain (CRD) have shown that Dectin-1A head groups form dimers when laminaritriose is present [57]. Additionally, size exclusion chromatography with multi-angle light scattering analysis has described Dectin-1 ligand-induced tetramer (or dimer-of-dimers) formation in solution [57, 58]. In line with these ideas, we sought to examine how ligand structure impacts signaling by determining the dimerization/oligomerization of full length Dectin-1A in living cell membranes.

To assess changing molecular proximity of Dectin-1 proteins, we employed Fluorescence Lifetime Imaging Microscopy for Forster Resonance Energy Transfer (FLIM-FRET). FRET based imaging capitalizes on close proximity of two proteins to visualize protein-protein interactions, including receptor dimerization and receptor-ligand complex formation [59]. FLIM characterizes the duration of a fluorophore’s excited state before returning to the ground state. The occurrence of FRET causes rapid quenching of donor fluorescence, so FRET can be determined by measuring the shortening of donor fluorescence lifetime when in proximity to acceptor. FLIM-FRET offers the opportunity of studying *in vivo* receptor interactions in a direct, spatially resolved manner. We examined Dectin-1A engagement using FLIM-FRET on HEK-293 cells co-expressing two fluorescent Dectin-1 constructs—N-terminally tagged Dectin-1A-mEmerald (donor) and Dectin-1A-mCherry (acceptor)—both having their fluorophores on the cytoplasmic face of the plasma membrane, and then stimulated the cells with soluble β-glucan. Analysis was conducted on the plasma membrane itself by masking out internal cellular compartments (Fig. 5A). FRET efficiency is a parameter that exhibits an inverse 6th power dependence upon donor-acceptor distance. Donor-acceptor percentage is simply the percentage of all donors that are involved in FRET interactions at a given time. These parameters were determined experimentally by performing bi-exponential curve fits to the observed frequency distribution of donor fluorophore lifetimes, wherein one exponential component represents the population of non-FRET monomeric donors and the other exponential component represents the performance of the donors that are involved in FRET interactions.

**Figure 5:**
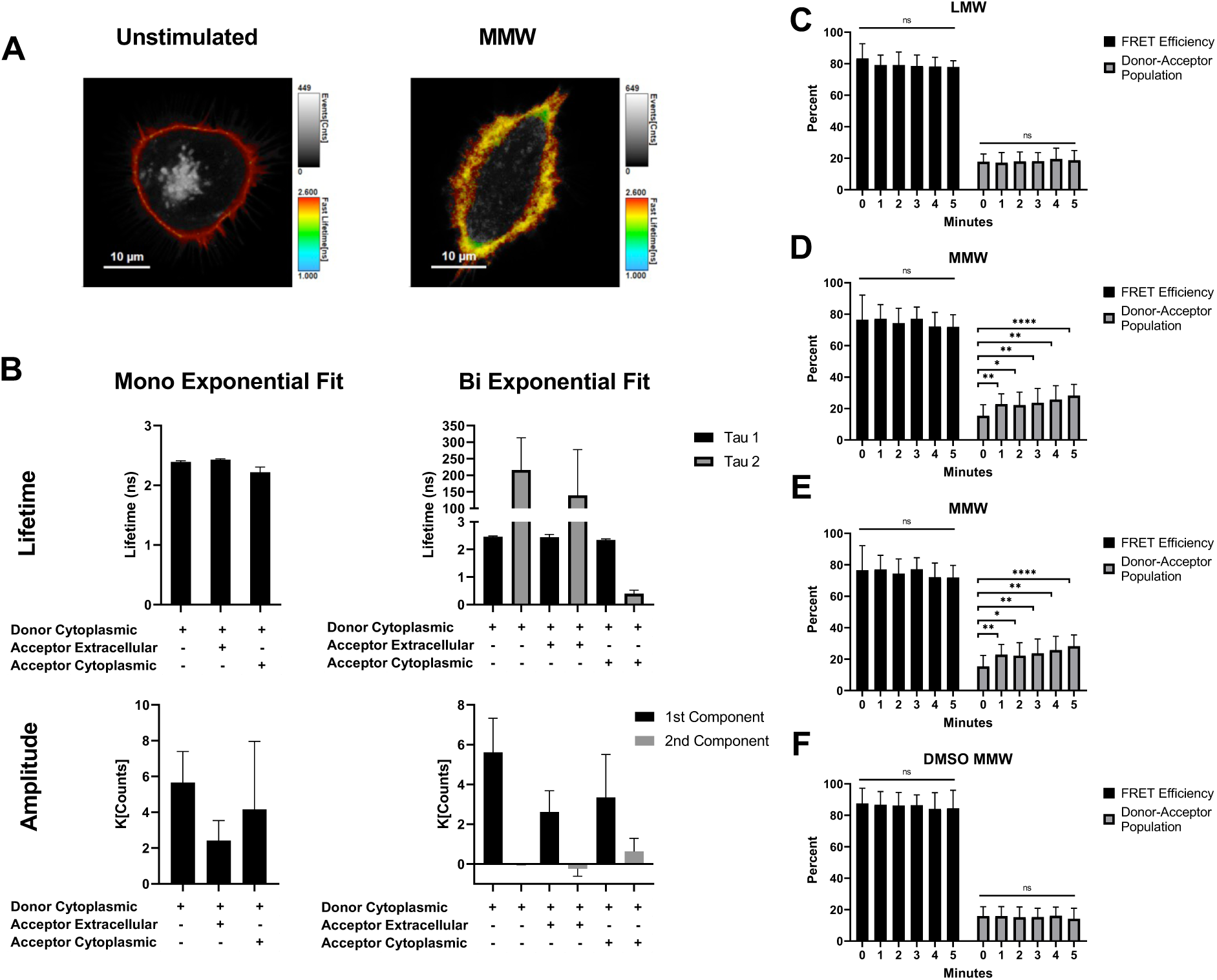
Highly structured β-glucans induce dimerization/oligomerization of Dectin-1A. (A) Representative average lifetime image of HEK-293 cell transfected with Dectin-1A-mEmerald or co-transfected with Dectin-1A-mEmerald and Dectin-1A-mCherry. The analysis was conducted on the plasma membrane by masking out internal cellular compartments on the images so only the masked plasma membrane signal used for analysis is shown in color. In these images, pixel hue indicates raw fluorescence lifetime over all populations observed and pixel intensity indicates total photon counts observed over all populations. (B) Lifetime and amplitudes of HEK-293 cells expressing donor only on the cytoplasmic face, donor expressed on the cytoplasmic side and acceptor on the extracellular side or donor and acceptor placed on the cytoplasmic side of the membrane. Fluorescence decay curves were mono- and bi-exponentially fit and individual fit components are shown. Data shown as mean ± SD (n = 15 cells). (C-F) FRET efficiency and Donor-Acceptor Population of cells stimulated with LMW (C), (D) MMW, (E) HMW, or (F) DMSO denatured MMW glucan. Data shown as mean ± SD (n = 15); One-way ANOVA with multiple comparisons by the Dunnett test, * p<0.05, **p<0.01, ***p<0.001.

First, we characterized the average lifetimes of unstimulated cells expressing several configurations of fluorescent protein tagged Dectin-1A: 1) receptor with donor tag only, 2) co-expression of separate donor and acceptor tagged receptors with tags placed on opposite sides of the plasma membrane (a negative control containing both fluorescent proteins but in a configuration that does not permit FRET), and 3) co-expression of separate donor and acceptor-tagged receptors with both tags in the cytoplasmic tail of the receptors (configuration to be used for experimental determination of receptor aggregation by FRET) (Fig. 5 B). The observed decay curves were analyzed by performing a mono-exponential and bi-exponential fit. For donor only and our negative control we observed a negative amplitude for the second component in the bi-exponential fit, indicating a mono-exponential fit was superior for these conditions. This was as anticipated since these conditions should have only a single, non-FRET donor signal. Additionally, the lifetimes of the second component in these controls unrealistically exceed >150 ns, further justifying a mono-exponential fit. Our results show that when acceptor is not present, we see the expected lifetime of 2.4 ns using a mono-exponential fit (Fig. 5B). Negative controls with donor and acceptor on opposite sides of the membrane yield similar lifetime values. Data from co-expression of cytosolic donor and acceptor-tagged receptor was fit bi-exponentially and the lifetime of both components is shown (Fig. 5B). We observed a decrease in the lifetime of the donor to 0.41 ns (FRET-involved second fit component of a bi-exponential fit) indicating that some basal level of intermolecular Dectin-1 close proximity interactions were being observed in unstimulated cells. Of course, the first fit component lifetime (non-FRET involved donors) remained at ∼2.4 ns, as expected from controls above. The existence of this basal FRET signal is interesting, and the potential sources and interpretation of this observation are further considered below. However, we first focused on assessing ligation-dependent changes in Dectin-1A’s molecular aggregation state as influenced by various glucans and measured by FRET. For the remainder of the FLIM-FRET experiments, we fixed the lifetime of the first component to 2.4 ns, and we showed the second components’ lifetime and amplitude values as percentages, FRET Efficiency and Donor-Acceptor Population respectively.

When we stimulate cells expressing Dectin-1A with donor and acceptor on the cytoplasmic face of the membrane using MMW (Fig. 5D) or HMW (Fig. 5E) glucan, we see a significant increase in the fraction of receptors undergoing FRET (Donor-Acceptor population) from 15% before stimulation to a maximum of 30% after 5 minutes of stimulation, with this trend starting at about one minute post-stimulation. However, there is not a significant change in FRET efficiency before and after stimulation. On the other hand, when we stimulate with LMW (Fig. 5C) or denatured MMW (Fig. 5F), we see no significant change in FRET efficiency or donor-acceptor population. We interpret the high and constant FRET efficiency for the population of receptors engaged in FRET interactions to mean that Dectin-1A in its aggregated state is in a close configuration (e.g., dimer or tetramer) that does not permit a wide range of separations between donor and acceptor tags, leading to a constant FRET efficiency for the donor population that does attain this FRET-capable configuration. However, the size of the population of receptors engaged in these close molecular aggregates does change as a result of stimulation with glucan. These results suggest that the highly structured soluble glucans allow for an increase in Dectin-1A dimerization or oligomerization to occur, which directly correlates with the amount of receptor activation and signaling observed.

In addition, to better characterize the aggregation states accessible to Dectin-1A, we conducted a Number and Brightness analysis (N&B) on HEK-293 cells expressing Dectin-1A-mEmerald. Previous research has described N&B more in depth [60–62]. Briefly, N&B analysis focuses on fluctuation of detected emission photons originating from fluorescent molecules that pass through a known observation volume. Statistics of fast fluctuations of the intensity at each pixel can be used to determine the number and intensity of the particles diffusing through the observation volume. For example, if the fluorescent proteins diffuse as a tetrameric protein, we expect to observe emission intensity fluctuation with four times more photons relative to a monomeric fluorescent protein diffusing through the excitation volume. Receptor aggregation was observed by stimulating these cells with soluble glucans. A brightness vs intensity 2D histogram of each pixel in a time series was developed and selection boxes were drawn to represent monomers (red box), dimers (green box) and oligomers (blue box) (greater than dimer; Fig. 6A). Dectin-1 aggregation state maps of representative untreated (Fig. 6B) and MMW-stimulated cells (Fig. 6C) were generated using this color scheme. Our results indicated that unstimulated and LMW stimulated cells contained a significantly higher amount of monomer pixels compared to MMW and HMW glucan treated cells (Fig. 6D). Inversely, we observed a significant increase in pixels with a dimer and oligomer brightness in cells stimulated with MMW and HMW compared to cells that were unstimulated or LMW stimulated (Fig. 6E, F). N&B analysis revealed that dimers of Dectin-1A account for the majority of aggregated state Dectin-1.

**Figure 6:**
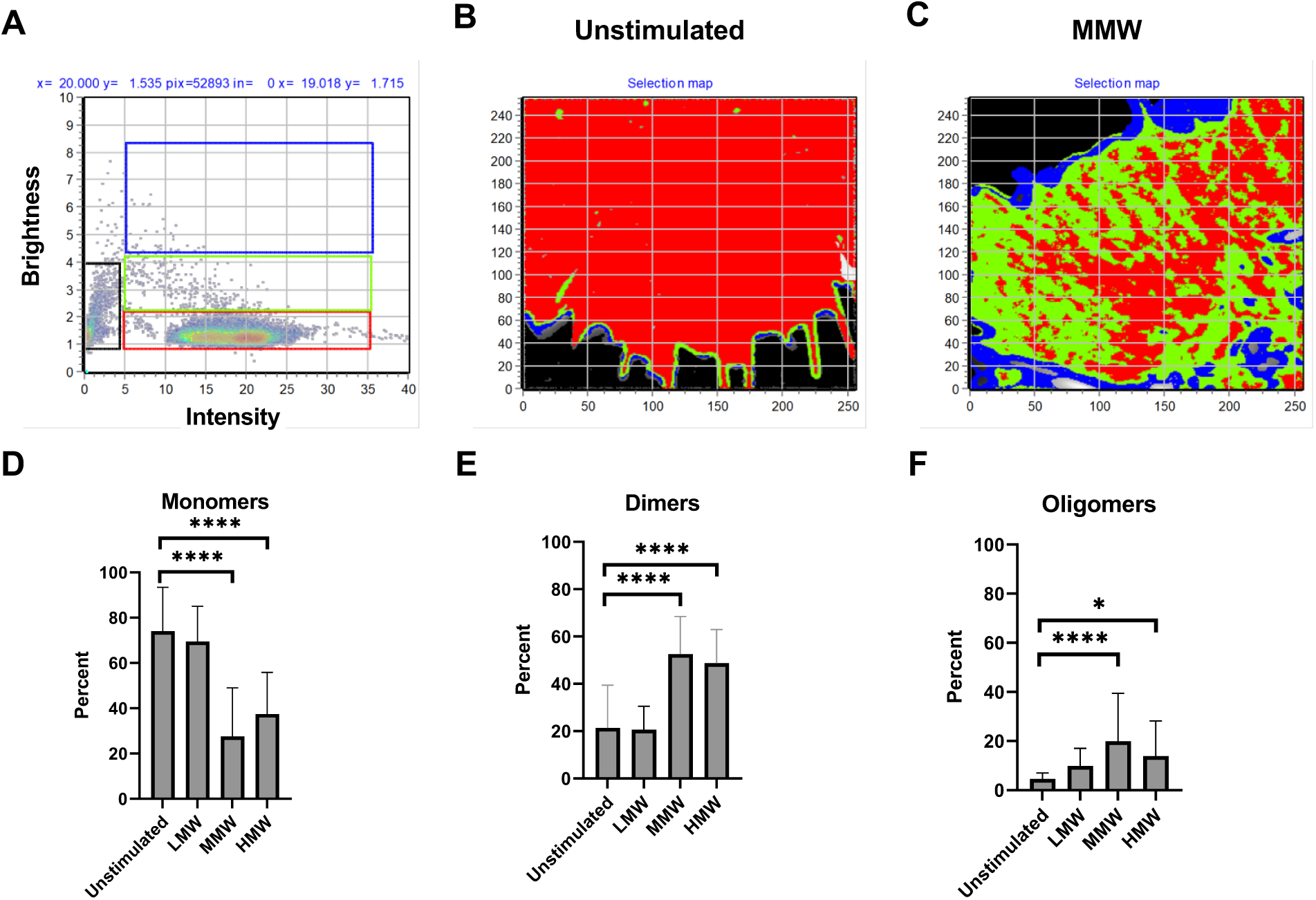
Number and Brightness analysis shows formation of small oligomeric states of Dectin-1A when stimulated with highly structured β-glucans. (A) Brightness vs intensity 2D histogram shows the selected pixels that contribute to the background (black), monomers (red), dimers (green), and oligomers (blue) in the image. (B) Representative selection map of HEK-293 cells expressing Dectin-1A-mEmerald shows receptor aggregation in unstimulated cells or those stimulated with (C) MMW β-glucan. Dectin-1 aggregation states are defined by colored boxes selected in the Brightness vs intensity histogram. (D,E,F) Percentage of (D) monomers, (E) dimers, and (F) oligomers in Dectin-1A-mEmerald receptors and receptor ligand complexes obtained from N&B analysis before or after stimulation. Data shown as mean ± SD (n = 30); One-way ANOVA with multiple comparisons by the Dunnett test, * p<0.05, **** p<0.00001.

### β-glucan induced Dectin-1A aggregates are below 15 nm in size

While FRET-based observations and N&B analysis clearly show the presence of ligand-induced, molecular-scale aggregation (e.g., dimerization) of Dectin-1A, these methods are not as well suited to discern the existence of larger scale aggregation of the receptor (e.g., clusters of tens to hundreds of receptors). We used direct Stochastic Optical Reconstruction Microscopy (dSTORM) coupled with Hierarchical Single-Emitter hypothesis Test (H-SET) analysis [13] to resolve aggregation of Dectin-1A before and after stimulation with MMW glucan. This localization super resolution microscopy technique accurately resolves objects from the diffraction limit (∼300 nm, the resolution limit of conventional fluorescence microscopy methods) or above, down to ∼15 nm (a typical resolution limit of dSTORM using our configuration). H-SET analysis detected sites of Dectin-1 labeling as “singlet” objects or “multiple” clustered objects. Multiple clustered objects are those with three or more resolvable individual Dectin-1 molecules. Singlet objects are those that appear to contain only a single, resolvable Dectin-1 labeling event, though it is possible that multiple Dectin-1 molecules in very close proximity (<15 nm separation) would be unresolvable and appear as a singlet object. We detected no significant change in the density of singlet objects or multiple object clusters before vs after MMW glucan stimulation (Fig. 7A,B; Supplemental Fig. 2). Consistent with a Dectin-1 distribution of predominantly monomers or low order oligomers (likely unresolvable by dSTORM), singlet exposures greatly outnumber multiple exposures on the cell wall surface (Fig. 7C). Localization number per multiple cluster object did not change with stimulation (Supplemental Fig. 2), suggesting no change in the number of receptors in this minority population of Dectin-1. In the context of the previous findings showing molecular aggregation at very small scales, potentially below the resolution limit of dSTORM, we concluded that dSTORM results indicated that the Dectin-1A aggregates formed upon glucan stimulation are quite small and remain below the resolution limit of dSTORM (<15 nm length scale). Because this places an upper bound on the size of ligand-induced Dectin-1 clusters and we can estimate that the CRD of Dectin-1 occupies an area approximately 25 nm^2^ (PDB: 2BPD; [57, 63]), we conclude that Dectin-1 aggregation after MMW glucan stimulation most likely involves collections of not more than ∼7 receptors.

**Figure 7:**
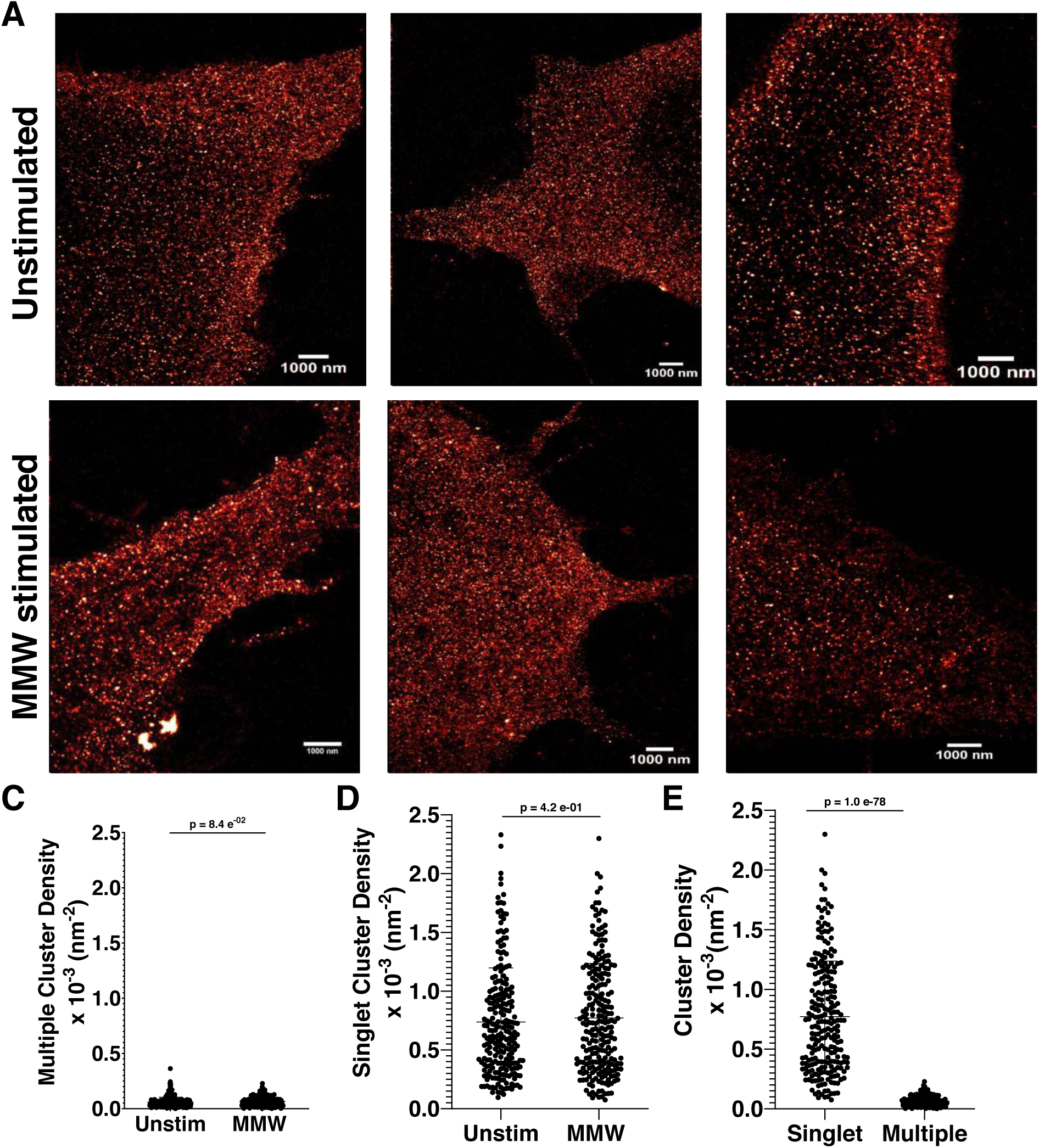
Dectin-1A does not form large scale aggregates when stimulated with highly structured β-glucan. (A) Representative immunofluorescence staining images of HEK-293 cells expressing Dectin-1A and either unstimulated or stimulated with MMW β-glucan. Cells were stained with a conjugated anti-Dectin-1-Alexa 647 antibody. (B) Multiple cluster density of dSTORM analysis of HEK-293 cells expressing Dectin-1A unstimulated or stimulated for 50 sec with MMW glucan. (C) Singlet cluster density of dSTORM analysis of HEK-293 cells expressing Dectin-1A unstimulated or stimulated for 50 sec with MMW glucan. (D) Cluster density of singlet and multiple exposure of Dectin-1A expressing HEK-293 cells treated for 50 sec with MMW glucan. Data shown as mean ± SD (n = 34) with significance assessed by Student’s T Test.

### Dectin-1A is predominantly monomeric in resting cell membranes

FRET-based measurements and N&B analysis reported that the large majority of Dectin-1 is distributed as monomers in unstimulated cells. However, a minority population of apparent close-proximity receptor states was observed in resting cells by both techniques. This may represent density-dependent, close-proximity interactions between Dectin-1 molecules driven by random collisional interactions, without necessarily requiring receptor oligomer formation. Alternatively, it is possible that a small fraction of Dectin-1 does form low order oligomers, even in the absence of glucan. It is difficult to conclusively distinguish between these alternative hypotheses using only the experimental results shown above. Therefore, we created a computational model of fully monomeric Dectin-1 undergoing Brownian 2D diffusion. If such a model could predict collisional FRET interactions at a level consistent with our FRET observations in resting cells, we would conclude that random collisional interactions are sufficient to explain basal FRET observed for Dectin-1 in this study. To accurately parameterize this model, we determined the receptor density of both donor and acceptor for HEK-293 cells coexpressing Dectin-1A mEmerald/mCherry by RICS analysis (Fig. 8A). Our results indicated on average Dectin-1 cotransfected cells contain 54.6% mEmerald and 45.4% mCherry (Fig. 8B). Therefore, our model was populated by equal proportions of donor and acceptor tagged Dectin-1 molecules. Their diffusion coefficients were parameterized using data from Fig. 4. The model calculated FRET rates for all donor-acceptor pairs within a specified maximum radial distance. The maximum radius for experimental FRET observation is limited by signal-to-noise ratio and other factors. To avoid simulating FRET measurements at experimentally unrealistic radii, we varied the model’s maximum donor-acceptor radial distance for FRET calculations in a range of 4-10 nm. We compared simulated and experimental FRET efficiencies and used maximum radii for simulations that yielded FRET efficiency in best agreement with observed FRET efficiency on resting cells, indicating comparable “sensitivity” of FRET detection in both. Our results show our experimental FRET efficiency values match model predictions closely at maximum radii between 5 and 6 nm (Fig. 8C). Simulations at these chosen parameters were then compared to experimental results with respect to the donor-acceptor population percent that they predicted.

**Figure 8.**
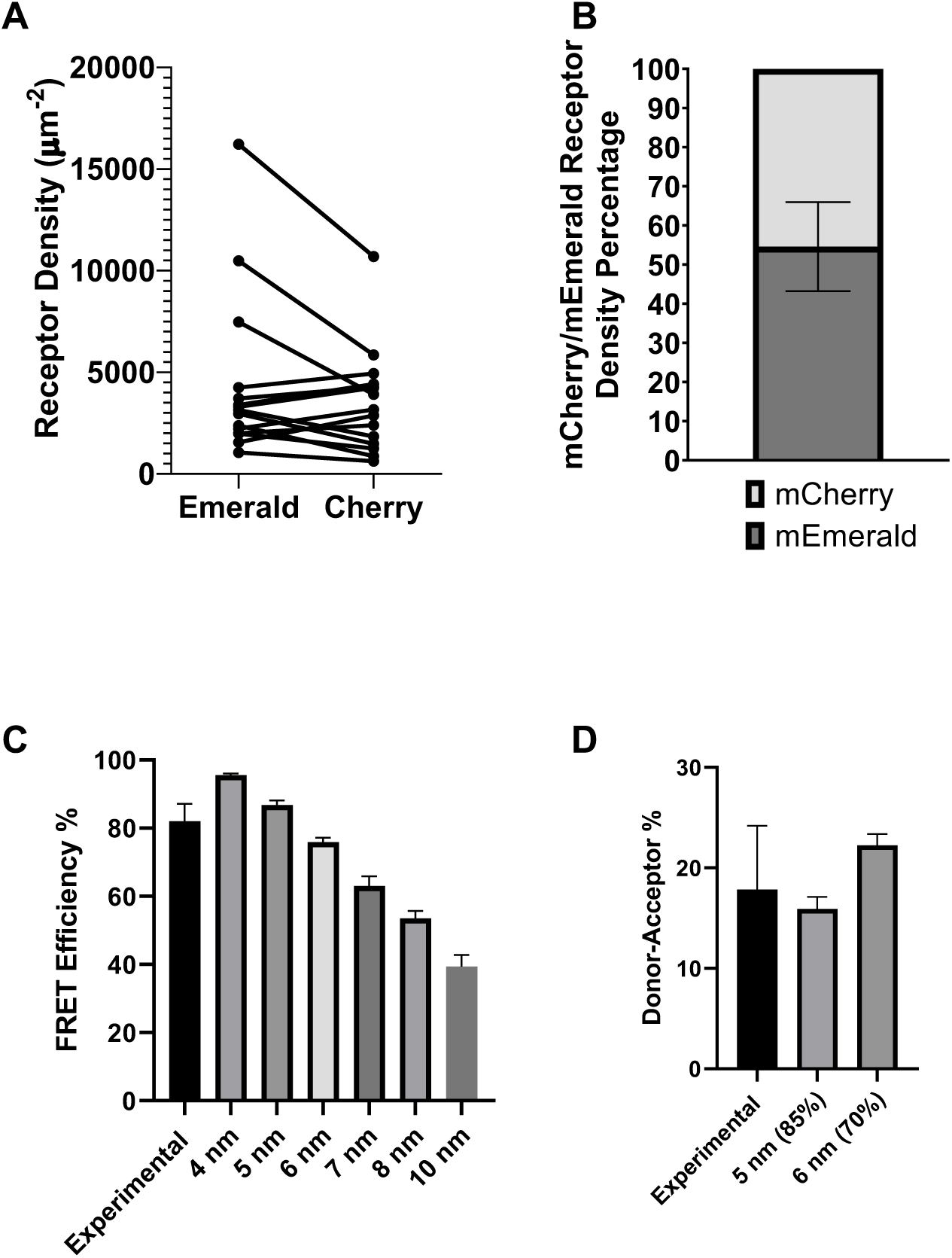
Computational FRET modeling of monomeric Dectin-1A can account for basal FRET observed in Dectin-1A expressing cells. (A) Receptor density of HEK-293 cells co-expressing Dectin-1A-mEmerald and -mCherry. Lines connect paired readings from single cells. (B) Ratio of mEmerald/mCherry expression in unstimulated HEK-293 cells. (C) FRET Efficiency of Dectin-1A donor/acceptor-containing computational models at different maximum intermolecular radial distances and experimental observations of FRET efficiency in resting cells for comparison. To facilitate comparison, model FRET Efficiencies are averaged over only the donors participating in non-zero FRET efficiency interactions with acceptors, and experimental FRET efficiency is derived from aggregation of all unstimulated (“0 minute”) cell data depicted in Fig. 5C-F. (D) Percentage of donors undergoing FRET (Donor-Acceptor Population) for experimentally observed populations in unstimulated cells and computational FRET modeling data at 5 and 6 nm maximum FRET radii. Experimental FRET donor-acceptor percentages were derived from aggregation of all unstimulated (“0 minute”) data in Fig. 5C-F. Data shown as mean ± SD (n = 15 independent simulations or experimental observations on cells, respectively); One-way ANOVA multiple comparisons Dunnett test.

We used the predictions of this computational model to test the hypothesis that random collisional FRET interactions of donors and acceptors is sufficient to explain the basal FLIM-FRET signal observed experimentally in resting cells. If this model, which incorporates only collisional interactions between donors and acceptors, predicts a percent of donors undergoing FRET interactions with acceptors that match the experimentally observed value, we would consider that collisional interactions alone are sufficient to explain the observed basal FRET signal. However, if the model predicts a value significantly below that experimentally observed, we would propose that a minor fraction of Dectin-1 molecules may participate in oligomeric aggregates on cell membranes, even in the absence of glucan. Using simulations with maximum radial values of 5 and 6 nm, our results indicate that the experimentally observed amount of Dectin-1 receptors dimerizing (Donor-Acceptor population) prior to stimulation match closely to our simulated results (Fig. 8D). This indicated that the FRET signal we observe prior to stimulation was attributable to random “collisional” interactions of Dectin-1A at the level of expression present in our experimental system.

### Dectin-1A dimer/oligomer and contact site formation is more efficient with cells incubated with *C. albicans* containing high glucan exposure

Throughout the article, we have focused on experiments using soluble glucans. In order to show Dectin-1 aggregation occurs during fungal pathogen recognition, we used a high (TRL035) and low (SC5314) β-glucan exposing *C. albicans* yeast and examined Dectin-1 aggregation occurring at the contact sites. TRL035 has been previously shown to have high glucan exposure compared to SC5314 [64]. In addition, our representative images show TRL035 forming a phagocytic cup more efficiently than SC5314 (Fig. 9A). Our results show that HEK-293 cells co-transfected with Dectin-1A-mEmerald and Dectin-1A-mCherry exhibit an increase in the proportion of aggregated-state receptors from approximately 15% in non-contact site membrane to about 25% in contact sites of high glucan exposing yeast TRL035 and *C. albicans* derived glucan particles.

**Figure 9:**
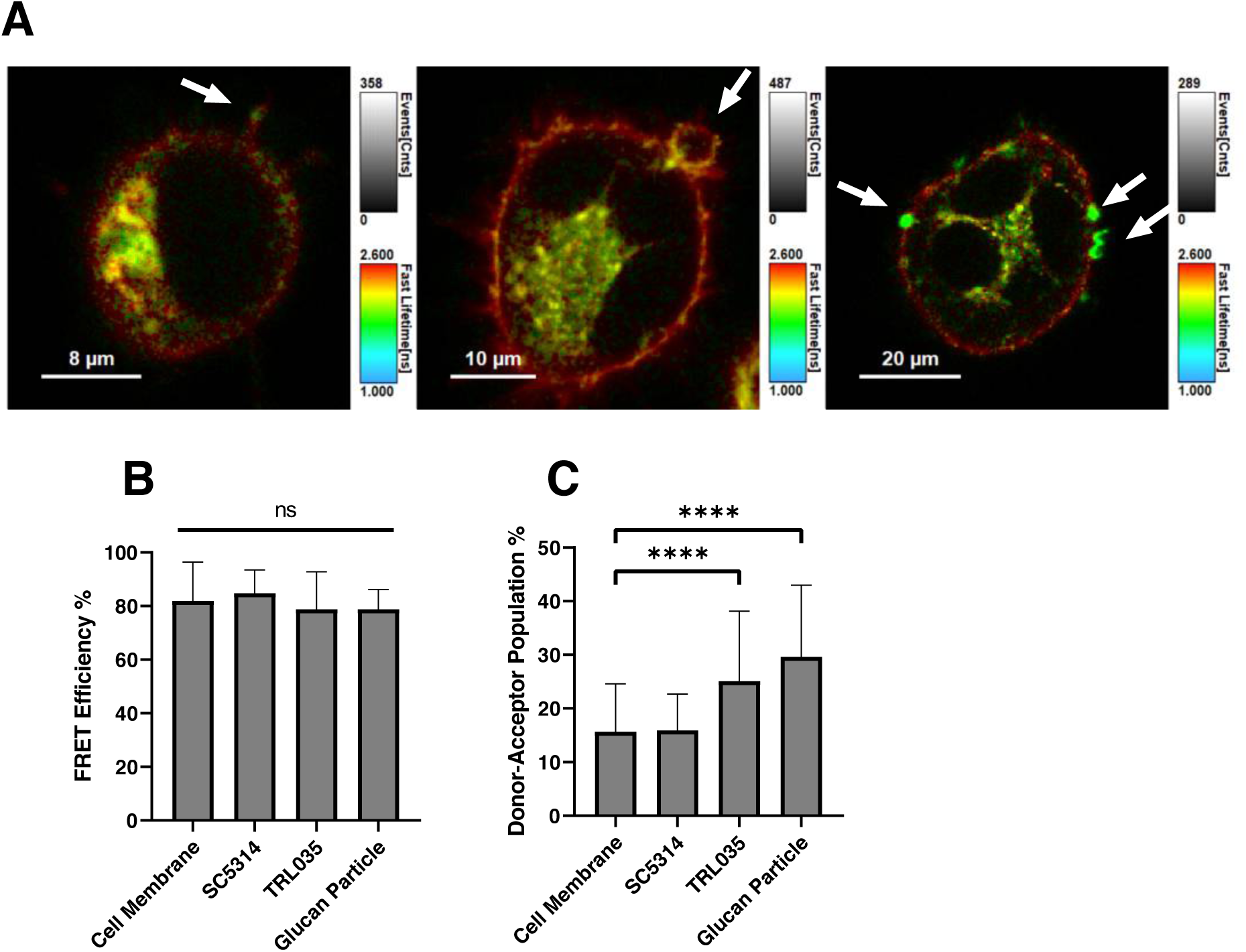
Dectin-1A forms more oligomers at fungal contact sites with high β-glucan exposure. (A) Representative lifetime images of HEK-293 cells co-transfected with Dectin-1A-mEmerald and Dectin-1A-mCherry incubated with SC5314 (left panel), TRL035 (middle panel) or particulate glucan (right panel). Transition from redder to greener pixels at contact sites is indicative of increased FRET interactions between Dectin-1 receptors. In these images, pixel hue indicates raw fluorescence lifetime over all populations observed and pixel intensity indicates total photon counts observed over all populations. (B) FRET efficiency and (C) donor-acceptor population of cell membranes with no fungal contact, SC5314 (low glucan exposure), TRL035 (high glucan exposure), or particulate glucan. Data shown as mean ± SD (n = 15); One-way ANOVA with multiple comparisons by the Dunnett test, **** p<0.00001.

Interestingly, we did not measure a significant increase in receptor aggregation between non-contact membranes and contact sites with low glucan exposing SC5314 (Fig. 9 B,C), which may indicate that the amount of aggregated Dectin-1 at SC5314 contacts was quite small and below the detection limit. Furthermore, we observed no significant difference in FRET efficiency between any conditions tested (Fig. 9B), similarly to our observations with soluble glucan. The significance and interpretation of these findings is further discussed below. These results suggest that the larger cell wall glucan exposure results in an increase of Dectin-1 in molecular aggregates with associated signaling, resulting in a more efficient recognition of yeast by the Dectin-1A receptor.

## Discussion

Our results demonstrate that the structure of β-glucan impacts receptor signaling by determining the membrane organization and molecular aggregation state of Dectin-1A. We showed that glucans with higher order structure are better able to activate Dectin-1A signaling. Upon activation by stimulatory soluble glucan, Dectin-1A enters aggregated states that contain dimers and higher order oligomers, but these appear to remain as small nanoscale domains containing relatively small numbers of receptors. Comparison of computational modeling and experimental FLIM-FRET data confirms that Dectin-1A exists in a monomeric state in resting cells. Further, monomeric receptors oligomerize and aggregate in a fashion that is dependent upon glucan higher order structure and correlated with the magnitude of membrane proximal signaling downstream of Dectin-1. Finally, we observed that a similar process of increasing Dectin-1 aggregation is seen at contact sites with yeast and fungal-derived particles, and that the amount of aggregated state Dectin-1A correlates with the degree of glucan exposure on the surface of the particle.

The prevalence of ensemble based studies of biological response to fungal glucans, using glucans from varying sources and with varying degrees of structural characterization, has complicated a thorough understanding of the impact of glucan physicochemical properties on their biological activity. Innate immunocytes are naturally exposed to β-glucan in both particulate and soluble forms. β-glucan is widely distributed on various fungal species as an insoluble component of the cell wall. The soluble form is produced when macrophages recognize fungal surfaces and release enzymes that degrade cell wall glucans [46, 47]. These soluble glucans are commonly found in circulation in the serum of patients with fungal infections [48, 49]. Low molecular weight glucans typically possess a random coil structure while increasing single or triple helical structure is generally seen as molecular weight increases [65]. In general, complexity of β-glucan correlates with immunostimulatory potency [17,27,29,65,66]. Mueller, et al indicated a correlation between glucan triple helical structure and binding affinity for receptors on human promonocytic cells, which is contrary to our finding of similar affinity across glucans tested herein [27]. Discrepancies in the impact of glucan structure on affinity may be due to different cell backgrounds, or more likely, to the fact that Mueller, et al used glucans from a wide range of sources, with complex differences in structure not merely limited to size or helical content. Our study used a carefully characterized series of fungal glucans to determine that Dectin-1A activation is specifically influenced by the degree of β-glucan triple helical structure, not merely through its affinity or size. Furthermore, the single-cell nature of our observations suggests that the biological response to glucans with strong helical structure (e.g., HMW glucan) seems bi-stable in nature, with response/non-response being correlated with glucan structure but the amplitude of calcium signal being similar in single cells, once successfully triggered.

Aggregate states of Dectin-1 relevant to signaling could exist in a pre-formed, ligation-independent state, be formed in a purely ligation-dependent manner, or a mixture of both models. We entertained the possibility of pre-formed aggregates because receptor oligomeric states exist in the basal state for some other C-type lectin receptors, such as DCSIGN, DNGR-1, and NKp80 [67–70]. However, Dectin-1A does not contain cysteine residues in its stalk region which are important in the dimerization of some other C-type lectin receptors (i.e., DNGR-1). Comparison of FRET data and computational modeling results demonstrated that any FRET activity observed in resting cells was explainable by expected levels of transient collisional donor-acceptor interactions in our cells. N&B analysis independently confirmed that the large majority of Dectin-1 is in monomeric states prior to stimulation. Therefore, we concluded that Dectin-1 is unlikely to undergo significant dimerization/oligomerization prior to ligation by glucan.

We proceeded to test the hypothesis Dectin-1 undergoes ligation-dependent aggregation. Specifically, we investigated whether glucan structure impacts signaling by modulating the frequency of Dectin-1A dimer/oligomer formation. Dectin-1 dimerization/oligomerization would create sites where Syk could be better recruited via interactions of its SH2 domains with the (hem)ITAM phosphorylated YXXL sequence in closely juxtaposed Dectin-1A cytosolic tails. In fact, this model is commonly cited in review literature in the field, but direct evidence in intact, live cells has been lacking [71, 72]. The plausibility of the Dectin-1 (hem)ITAM aggregation model is suggested by the fact that another C-type lectin receptor, CLEC-2, forms a minimal signaling unit composed of a phosphorylated dimer, enabling recruitment of a single molecule of Syk [44]. Our FLIM-FRET and N&B results reveal that Dectin-1A enters a state of greater molecular aggregation when stimulated with β-glucans, and that the degree of glucan helical structure correlates with its ability to induce Dectin-1 aggregation. FRET and N&B are very sensitive methods to identify receptor aggregation on the scale of small oligomers, but these methods are limited in their ability to distinguish such small aggregates from the formation of larger receptor nanodomains. dSTORM failed to detect ligand inducible Dectin-1 nanodomains on a length scale of ≥15 nm, suggesting that Dectin-1 aggregation events are limited to small collections of ≤7 receptors. Our core observation of ligation-inducible Dectin-1 aggregation is directly in line with previous crystallographic studies that show monomeric Dectin-1A CRD in the absence of glucan but able to form dimeric complexes in the presence of β-glucan [57]. In addition, solution biophysical studies have shown a ligand-induced cooperative formation of Dectin-1 CRD tetramers (or dimers of dimers) [57, 58]. However, these studies were performed with truncated receptor ectodomain proteins outside the context of living cell membranes, so our findings better establish and define the relevance of ligation-dependent Dectin-1 aggregation in a more physiologically realistic context. Nanoscale glucan exposures on *Candida* cell wall surfaces may be important determinants of the degree of Dectin-1 aggregation at host-pathogen contact sites [13, 64]. Dectin-1 is recruited to the “phagocytic synapse” between innate immunocytes and fungal particles. Here, Dectin-1A encounters fungal glucan and initiates signaling [73]. We have previously described that *C. albicans* TRL035 exhibits larger glucan nanoexposures than *C. albicans* SC5314 [64]. Consistent with the presence of larger glucan exposures, we observed greater Dectin-1 aggregation at FRET contact sites with TLR035, relative to SC5314. Cell wall glucan nanoexposures are larger (∼20-200 nm) than the Dectin-1 aggregates generated by soluble glucans in the present work. Therefore, future studies could productively investigate the role of glucan nanoexposures in stabilizing larger aggregated collections of engaged Dectin-1, and the potential dependence of cellular activation on the scale of Dectin-1 aggregation at sites of cell wall glucan nanoexposures.

These and other studies improve our physical understanding of host-*Candida* interaction and highlight the exquisite sensitivity of the Dectin-1 system that drives innate immune fungal recognition. Our previous optical nanoscopy studies of *Candida* cell wall surfaces estimated multivalently-engaging glucan exposure site density and area (per exposure site) as follows: SC5314—1 µm^-2^ density, 6.61×10^-4^ µm^2^ area; TRL035—4 µm^-2^ density, 9.62×10^-4^ µm^2^ area [64]. The total area of contact sites between *C. albicans* and human immature dendritic cells is ∼10 µm^2^ [74]. Finally, we estimate (see above) that one Dectin-1 CRD occupies a footprint of ∼25 nm^2^. From these figures, we calculate that a typical phagocytic synapse would contain a maximum of 264 multivalently engaged Dectin-1 proteins for *C. albicans* SC5314, and maximum 385 multivalently engaged Dectin-1 for TRL035 (at total ligand engagement). Based on our reported Dectin-1 density (Fig. 4), the contact sites we measured would contain ∼46000 total Dectin-1 proteins. So, the Dectin-1 system is able to drive signaling responses when, at most, only a few hundred receptors, corresponding to less than 1% of the total contact site resident Dectin-1 proteins, are aggregated in the contact. These results and estimates suggest that fungal recognition requires the Dectin-1 system to engage in a search for rare sites of multivalent interaction with glucan. Signal initiation must be sensitive to activation of relatively small numbers of Dectin-1 proteins. In the future, it will be important to achieve a better understanding of Dectin-1’s collaboration with other anti-fungal receptors (e.g., DC-SIGN and CD206). Such receptors may be important for building and stabilizing a fungal contact that can effectively promote Dectin-1’s ability to search for and find its rare sites of glucan exposure.

Overall, these findings indicate that β-glucan structure is required for Dectin-1A to undergo Syk-dependent signaling. Here we provide evidence in support of a model in which highly structured glucans induce stable dimerization and/or oligomerization of the receptor. This allows their (hem)ITAM domains to become close enough for a sufficient period of time to allow for the activation of Syk, leading to further signaling cascades. Greater understanding of receptor activation is required to better understand the role of Dectin-1A and its agonists as a potential way forward for adjuvant and immunotherapy development. Furthermore, given the worldwide burden of candidiasis, further experimentation is required to better understand the role of Dectin-1A in recognition of these pathogens.

## Materials and Methods

### Cell Culture

The HEK-293 (ATCC, #CRL-1573) cell line was maintained in Dulbecco’s Minimum Essential Medium supplemented with 10% Fetal Bovine Serum (FBS), 1% penicillin/strepromycin, 2mM L-glutamine, 11mM sodium pyruvate, and 1% HEPES. Cells were grown in an incubator at 37°C at 5% CO_2_ and saturating humidity. Cells were maintained at 37°C, 5% CO_2_, and 75% relative humidity during imaging.

### Plasmids and Transfection of Dectin-1 Constructs

Emerald-Dectin1A-N-10(Addgene plasmid, #56291), Emerald-Dectin1A-C-10 (Addgene plasmid # 54057), mCherry-Dectin1A-C-10 (Addgene plasmid # 55025), and mCherry-Dectin1A-N-10 (Addgene plasmid # 55026) was a gift from Michael Davidson. pUNO1-hDectin-1A (Invivogen) was stably transfected into HEK-293 cells for use in our calcium studies. Stable transfection of mEmerald-Dectin-1A was used for Syk immunoblotting experiments. To generate stable lines, HEK-293 cells expressing either mEmerald-Dectin-1A or pUNO-hDectin-1A were selected using Geneticin (G418 Sulfate) (Thermo-Fischer, #10131035) at 400 μg/ml or Blasticidin (Santa Cruz Biotechnology, #SC-495389) at 20 µg/ml, respectively, for 2 weeks.

All other experiments involving exogenous protein expression used transient transfection. Transient transfection with plasmids was performed using standard manufacturer protocols with Fugene 6 (Promega, #E2691).

### Fungal Growth/Preparation

*C. albicans* SC5314 (ATCC, MYA-2876) or TRL035 yeast cells were grown from glycerol stock, stored at −80°C. Samples were grown in YPD, for 16 h at 37°C in an orbital shaker at 250 rpm to mid log phase. Following a 3-minute centrifugation at 6000 rpm, the supernatant was removed, and the cells were resuspended in 4% paraformaldehyde and sterile phosphate-buffered saline (PBS) for 15 minutes. The cells were centrifuged and washed with sterile PBS three times. The cell concentration was then determined using a disposable hemocytometer (C-Chip; Bulldog Bio catalog no. DHC-N01). 3.5 x10^6^ cells were resuspended in 1 ml of PBS. 100 µl of the solution was added to HEK-293 cells in 35 mm dishes 15 minutes prior to imaging.

### Glucan Particles

Glucan microparticles were prepared from lyophilized *C. albicans* SC5314 yeast derived from stationary phase culture in YPD. Dry yeast were extracted thrice in boiling 0.75N NaOH (15 min), then residue was extracted thrice in boiling 2N H_3_PO_4_ (15 min), then residue was extracted thrice in boiling acidic ethanol (1% v/v H_3_PO_4_ in ethanol; 15 min), and residue slurry was adjusted to neutral pH and washed thrice with ultrapure water. Pyrogen free reagent and glassware was used throughout preparation, and particles were stored at 4°C in sterile, pyrogen free water.

### Soluble Glucan Chromatographic Analysis

Low (LMW, 11 kDa), medium (MMW, 145 kDa), and high (HMW, 450 kDa) molecular weight β-(1,3;1-6)-glucan extracted from *S. cerevisiae* cell wall was obtained from ImmunoResearch Inc. (Eagan, MN). The molecular weight was assessed by gel permeation chromatography (GPC) and multi-angle light scattering (MALS). Samples (100 μg) were injected and eluted with a mobile phase of 0.15 M sodium chloride containing 0.02% sodium azide at a flow rate of 0.5 mL/min using two Waters Ultrahydrogel 500 columns and one Waters Ultrahydrogel 250 column connected serially. The samples were run with the column temperature at 18°C. The Mw was calculated using Wyatt Astra software using data resulting from measurements of the angular variation of scattered light using the MALS detector coupled with the concentration measured by the refractive index signal.

### Soluble Glucan Linkage Analysis

Desalted and lyophilized samples of the fractions were dissolved in dimethylsulfoxide (DMSO) and treated with NaOH and methyl iodide to methylate all free hydroxyl groups [75]. The methylated material was purified by extraction with dichloromethane and washing with water. The purified material was then hydrolyzed with trifluoroacetic acid, the reducing ends of the resulting sugars were reduced with NaBD_4_, and then the resulting free hydroxyl groups were acetylated with acetic anhydride. The mixture of partially methylated alditol acetates was analyzed by gas chromatography. Each derivative corresponding to a particular linkage has been identified by a characteristic retention time and mass spectrum using a mass detector. The relative amount of each derivative was measured by gas chromatography with flame ionization detection. The areas obtained for each observed peak were used to calculate the relative amounts of each type of linkage found in the sample (Table 1). The 3,6-linked residues represent branch points.

**Table 1.**
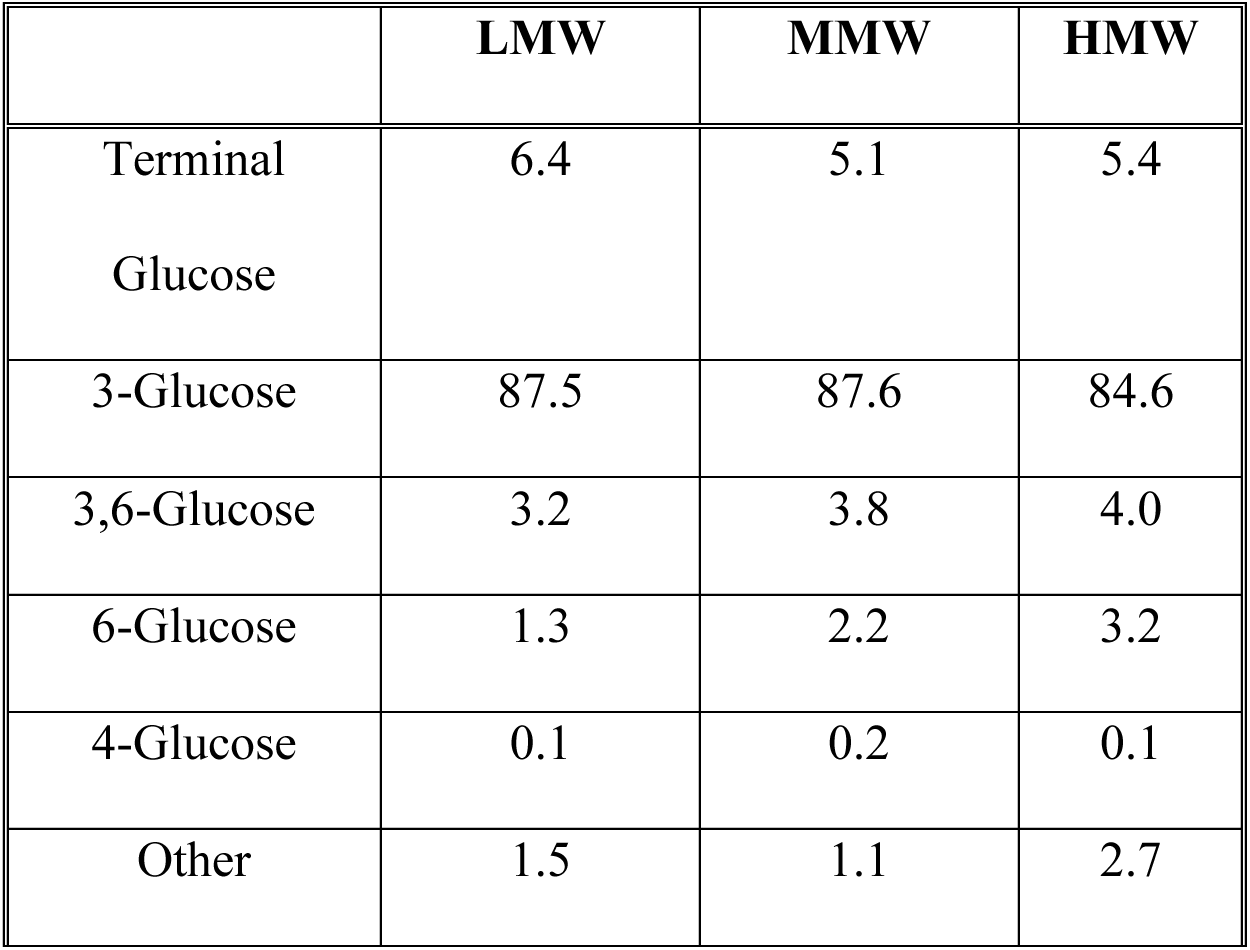

|  | LMW | MMW | HMW |
| --- | --- | --- | --- |
| Terminal<br>Glucose | 6.4 | 5.1 | 5.4 |
| 3-Glucose | 87.5 | 87.6 | 84.6 |
| 3,6-Glucose | 3.2 | 3.8 | 4.0 |
| 6-Glucose | 1.3 | 2.2 | 3.2 |
| 4-Glucose | 0.1 | 0.2 | 0.1 |
| Other | 1.5 | 1.1 | 2.7 |

### ^1^H NMR Spectroscopy

The samples were dissolved in DMSO-d6/D2O (6:1 by volume) at 100°C for 1 h. ^1^HNMR Spectra were recorded at the University of Minnesota Department of Chemistry NMR lab on a Varian UNITYplus-300 spectrometer at 80°C. The spectra were collected at 300 MHz with 32 scans, a relaxation delay of 1.5 seconds, a pulse of 45°, an acquisition time of 2.0 seconds, and a spectral width of 5999 Hz. Table 2 provides ^1^H NMR chemical shifts in all three glucans used in this study as well as literature values [50]. These data, taken together with other characterization methods used, do confirm that the structure of the polysaccharides used in this study conforms to expected results from fungal cell wall glucans (Table 2, Supplemental Fig. 3).

**Table 2.**
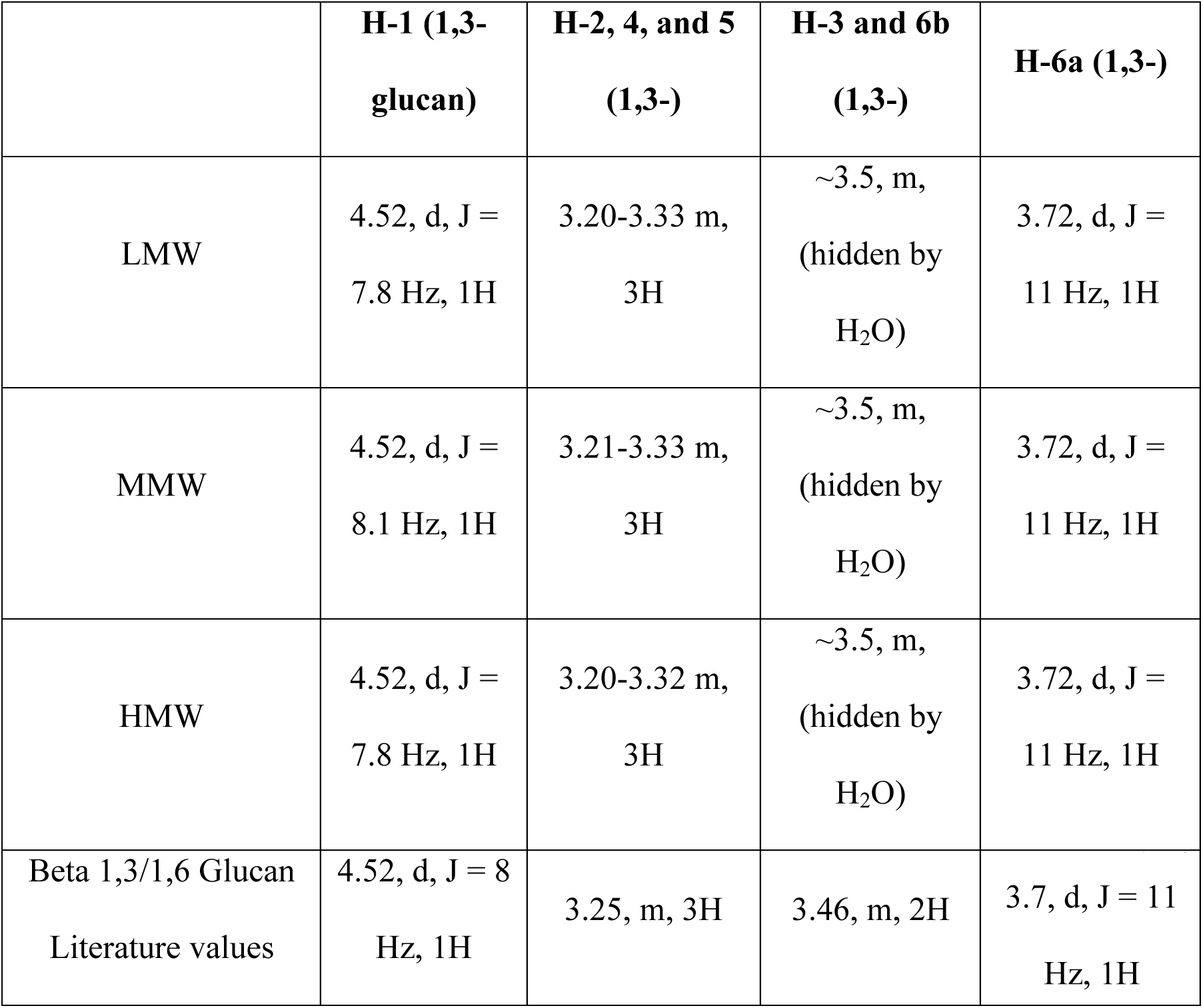

### Microscopy and Image Analysis (Calcium Imaging & RICS/N&B)

Confocal images were obtained on an Olympus FV1000 laser scanning confocal microscope (Olympus, Center Valley, PA) built around an IX81 inverted microscope. A 10x objective lens (0.40 NA) or a super corrected 60X oil objective lens (1.40 NA), Plan-Apochromat objective lens was used for imaging. Samples were excited with a 20mW, 473 nm diode laser and a 20 mW, 635 nm diode laser. These lines were reflected to the specimen by a 405/473/559/635 multi-edge main dichroic element followed by bandpass emission filters in front of 2 independent High sensitivity GaAsP PMT detectors (HSD1/2). Specifically, the emission light passed by the main dichroic was directed to our first detector (HSD1) via reflection from a SDM560 filter cube and passage through a BA490-540 nm bandpass filter. Our second detector (HSD2) received light passed by the SDM560 filter cube and routed through a BA575-675 nm bandpass filter.

### Calcium Imaging

HEK-293 cells expressing Dectin-1A were plated at 40,000 cells in a 35 mm (MatTEK dishes) 24 hours prior to imaging. These cells were loaded with Fluo-4 and Cell Mask Deep Red (CMDR) at equimolar concentrations of 1 µM in 2 ml of media for one hour then washed before imaging. Cell Tracker Deep Red was used as a cell cytosolic volume control to account for cytosolic changes from cell contraction that occurs during stimulation. For Syk inhibition, plates were pre-treated with 250 nM of Syk Inhibitor (Calbiochem, #574711) for 30 min under normal growth environmental conditions. Images were taken at a resolution of 256 x 256 with a dwell time of 2 µs on a 10x objective lens (0.40 NA). A 20 mW, 473 nm diode laser operated at 4% power and CMDR was excited with a 20 mW, 635 nm diode laser operated at 4% power. Fluorescence of Fluo-4 was collected by a cooled GaAsP PMT set to 700V, gain 1X and offset of 0%. CMDR signal was collected by a cooled GaAsP PMT detector set to 700V, gain 1X and offset of 0%. 30 frames prior to stimulation were used to set the basal fluorescence of the fluo-4 dye. After stimulation with 100 µl of 10µg/ml of glucan, cells were imaged for 100 frames. To assess changes in intracellular calcium concentration, we measured the ratio of Fluo-4/CMDR intensity in order to correct for any variations in cytoplasmic volume within the confocal section across the field. This ratio was normalized to 1.0 based on mean pre-stimulation values (30 frames) and changes in calcium influx were measured as fold change of this normalized ratio (MFI fold change). For MMW denaturation experiments, soluble β-glucans were weighed and resuspended in reverse osmosis purified H_2_O. In order to denature medium molecular weight glucan, we incubated MMW in DMSO or 1M NaOH. To renature the glucan from DMSO, we placed denatured MMW into Slide-A-Lyzer Dialysis Cassettes (Thermofisher, #66203) of a molecular weight cut-off of 2,000 Da and dialyzed against reverse osmosis purified H_2_O for 24 hours. To renature glucan from 1M NaOH, the solution was neutralized using 1M HCl.

### Protein isolation and immunoblotting

HEK-293 cells stably expressing mEmerald-Dectin-1A were seeded at 5 x 10^5^ in 6-well plates 24 hours prior to the experiment. Cells were stimulated with Low, Medium and High molecular weight β-glucan (1 mg/ml) for 5 minutes, then lysed. Cells were extracted in 1X lysis buffer (43.9 mM HEPES, pH 7.5; 131.7mM NaCl; 1.1% Triton X-100; 8.8% glycerol; 1x protease inhibitor cocktail; 1mM PMSF; 1mM EGTA). Samples were centrifuged at 12,000 x *g* for 20 min at 4°C and supernatants transferred to fresh tubes. Protein concentrations were determined by Bradford assay (Bio-Rad Protein Reagent). NuPAGE LDS sample buffer (4X) with NuPAGE Sample Reducing agent (10X) was added to samples (1X final concentration). Total proteins (typically 20-50 µg) were subjected to 4-12% sodium dodecyl sulfate-polyacrylamide gel electrophoresis (SDS-PAGE). Proteins were transferred to Immobilon-FL PVDF transfer membrane (Millipore Sigma) using NuPAGE transfer buffer. Membranes were blocked with bovine serum albumin in Tris-buffered saline-Tween-20 (TBS-T; 20 mM Tris, 137 mM NaCl, 0.1% Tween-20) and incubated with primary antibodies overnight at 4°C. Antibodies purchased from Cell Signaling: Rabbit mAb for p-SYK (Tyr525/526) and β-Actin (13E5), and mouse mAb Syk (4D10) were used according to manufacturer’s recommendations (1:1000). HRP-conjugated anti-mouse and anti-rabbit secondary antibodies (Cell Signaling or GE Healthcare) were used at a 1:10,000 dilution. Blots were visualized on a Li-Cor Odyssey FC imaging system and analyzed with Image Studio.

### Biolayer interferometry

Advanced Kinetics Biolayer interferometry experiments were conducted using the Personal Assay BLItz System. Anti-human IgG Fc Capture (AHC) Biosensors tips were initially loaded with Dectin-1A:FC fusion protein (Invivogen, #fc-hdec1a) at 13 ug/ml. Binding kinetics were obtained for LMW, MMW, HMW, MMW (denatured) and MMW (renatured) at 0, 10, 50, 100, and 250 nM in triplicate. A global fitting was performed on the curves obtained using the BLItz software.

### Congo Red Spectroscopic Assay

A BioTek EON Multiwell Spectrophotometer was used to analyze Congo red absorbances. A solution of 8.8 µM Congo Red, 0M-1M NaOH solution (1 M, 0.75 M, 0.5 M, 0.25 M, 0.1 M, 0.075 M, 0.05 M, 0.025 M, 0.001 M, 0 M) and LMW, MMW, or HMW β-glucans at 1 mg/ml were analyzed for the denaturation experiments. For the renaturation experiments, 1 mg/ml of β-glucan was denatured at 1M solution then renatured through neutralization with HCl 24 hrs prior to readings. DMSO denaturation conditions involved DMSO in water at 0%, 5% and 10%. For the DMSO renaturation experiments, DMSO was removed by dialysis (see above) prior to spectrophotometer readings. Absorbance readings were taken at 400-700 nm with 1 nm steps. All experiments were conducted in technical triplication across three independent experimental replicates.

### Fluorescence Lifetime Imaging Microscopy and Förster Resonance Energy Transfer Measurement

HEK-293 cells were plated at 25,000 cells in a 35 mm (MatTEK dishes) 48h prior to imaging. Cells were transfected with mEmerald-Dectin1A-N-10 and mCherry-Dectin1A-N-10 24 hrs prior to imaging. FLIM-FRET images were obtained using a Leica DMi8 inverted microscope. A Leica Harmonic Compound PL apochromatic CS2 63X water objective with a correction collar (1.2 NA) was used for imaging. A tunable & pulsed White Light Laser (470 - 670 nm) was operated at 80 MHz at 3% laser power using a 488 nm notch filter to excite our sample. A scan speed of 200 lines/sec and a 256 x 256 pixel resolution (full field of view) was used for data acquisition. Two hybrid detectors collected at photons at (512-540 nm) and (650-700 nm) respectively on the counting mode setting. Temperature was kept at 37 °C using a Tokai Hit Stage Top Incubator for Live Cell Imaging. Lifetime images were collected using a Pico Harp 300 Fluorescence Lifetime Microscopy Time-Correlated Single Photon Counting (TCSPC) system. For our glucan stimulated cells, prior to stimulation 23 frames were collected. 230 frames were taken immediately after stimulated with β-glucans at a concentration of 10 µg/ml. Analysis was conducted on minute time points (1-5 minutes) by averaging ten frames from one minute intervals. For yeast contact site imaging studies, 3.5 x10^6^ fixed yeast cells were resuspended in 1 ml of PBS. 100 µl of the solution was added to HEK-293 cells in 35 mm dishes 15 minutes prior to imaging. 23 frames were collected per cell. Images were collected at a maximum of 45 minutes after the addition of yeast per plate. Analysis was conducted on the plasma membrane by masking out internal cellular compartments on the images. For our fungal contact site studies, analysis was conducted on the plasma membrane that was in contact with the fungus and a separate masking for plasma membrane that was not in contact with any yeast. A bi-exponential fit was performed to the decay curve. For donor only as well as donor and acceptor on opposite sides of the plasma membrane (negative control), the decay curve indicated a negative amplitude for one of the components, thus indicating a mono-exponential decay. Therefore, decay curves from these samples were analyzed using a mono-exponential fit. For cells with donor-acceptor on the cytosolic tail, data was fit to a bi-exponential decay with the first lifetime component being locked at the donor only lifetime of 2.4 ns (τD). Lifetime values of the second component (τDA) of the decay curve were used to calculate FRET efficiency using the equation: 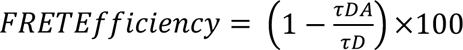. To determine the fraction of receptors undergoing a FRET process (Donor-Acceptor Population), the amplitude ratio between the first component (AmpD) and the second component (AmpDA) from the bi-exponential decay curve fit was calculated according to the following formula: 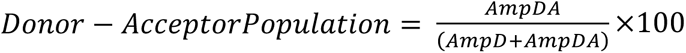.

### Raster Image Correlation Analysis/Number and Brightness

Protocols on RICS and N&B analysis have been previously described in more depth and analysis of diffusion coefficient and receptor density were performed using SimFCS software according to these previously published procedures [54, 76]. HEK-293 cells expressing Emerald-Dectin1A-C-10 were plated at 40,000 cells in a 35 mm glass bottom MatTEK dishes 24h prior to imaging using equipment described in “Microscopy and Image Analysis” section above. Measurements were performed at the membrane facing the glass coverslip. Images were collected at 256 x 256 pixel resolution on a 60× 1.4 NA oil immersion objective lens with a scanning zoom of 16.4X (0.050 µm vertical and horizontal center to center distance between resulting image pixels). Data was collected using a GaAsP PMT detector operated in photon counting mode. The 473 nm diode laser operated at 0.1% power was used in these images. The Point Spread Function (PSF) radial beam waist was estimated using 192 nM EFGP in solution and setting the diffusion coefficient to 90 µm^2^/s. Under these conditions the beam waist was determined to be 0.21 µm. Immobile features were removed using a 4-frame moving average subtraction. Cells were stimulated with a final concentration of 1 µg/ml of glucan. After stimulation, 200 frames were collected at a pixel dwell time of 4 µs/pixel (line scan of 1.096 ms) with a pinhole size of 100 µm. All cell body pixels were used for analysis of N&B data from cells.

Images collected were also used for our Numbers and Brightness analysis; we used 192 nM EGFP in solution and purified mEmerald-Dectin-1 protein to set the average brightness of our monomeric protein (Supplemental Fig. 4). Furthermore, the S-factor was calculated using the background image. We divided each brightness distribution by monomeric, dimeric, and oligomeric sections according to previous research [77]. The cursors were scaled quadratically and centered at B values of 1.3, 1.6, and 2.35 for monomers, dimers, and oligomers respectively.

### dSTORM

HEK-293 cells, were grown on cleaned and Poly-L-Lysine (0.1 mg/ml) coated coverslips (∼5 x 10^4^ cells/coverslip) within wells of a six-well plate at 37°C 24hrs prior to the experiment. The cells were then treated with MMW glucan at 1 µg/ml for 50 seconds. The cells were then fixed with paraformaldehyde (PFA; 4%) for 5 minutes at 37 °C followed by three washes of PBS.

Data acquisition was on an Olympus IX-71 microscope equipped with an objective based TIRF illuminator using an oil-immersion objective (PlanApo N, 150×/1.45 NA; Olympus) in an oblique illumination configuration. Sample excitation was done using a 637nm laser (Thorlabs, laser diode HL63133DG), with custom-built collimation optics [13]. To minimize the drift that occurred during data acquisition, a self-registration algorithm was implemented [78, 79].

The Dectin-1A nanodomain density by glucan exposure engagement was quantified by super resolution imaging and analyzed using H-SET as a clustering algorithm in MATLAB [13]. The data for 34 cells for each condition were run through the first pass of H-SET to collapse multiple observations of the blinking fluorophores into single estimates of the true fluorophore locations [13]. The second H-SET pass determined clustering using the DBSCAN algorithm [80] which depends on the two parameters. (minPts), that is, the minimum number of objects composing a multi-cluster, and the maximum distance between the objects within a multi-cluster (epsilon). We optimized these parameters, defining them as 3 and 27 nm, respectively, according to optimization procedures previously described [64].

### Collisional FRET Simulation with Monomeric Dectin-1

For this simulation, first a number of particles were generated based on the number of fluorophore molecules on the membrane in a specific membrane area on the real cell. As the experiments have shown that the ratio of donors to acceptors is roughly 1:1, in our model 50% of the particles were donors while the rest were acceptors. The absolute number of donor and acceptor molecules was based on the experimentally determined Dectin-1A membrane density presented in Fig. 8.

The initial location of each particle in the simulation space was defined by drawing random numbers from a uniform distribution. Throughout the simulation, particle movement was modeled by using a random walk process. The distance each particle moved at each time point was determined by the diffusion coefficient that was experimentally determined and reported in Fig. 4.

The simulation space included monomer molecules as donors and acceptors. The size of this simulation space corresponded to an experimental area equivalent to 0.16 µm^2^ of the cell membrane. The total duration of the simulation was equivalent to the total amount of time required to acquire data from 5 pixels of an experimental FLIM FRET observation (equivalent to 24 μs of total data acquisition time).

To simulate a TCSPC FLIM experiment, each simulation run included 2000 sequential excitation pulses, followed by a window of simulated fluorescence decay observation with a length of 12 ns (0.1ns time resolution). At the start of each pulse, 30% of the donors were selected to act as excited particles, which corresponded with fractional donor excitation observed under our FLIM experimental conditions (Supplemental Fig. 5). Note that we consider that this value represents an upper bound on % donors excited under experimental conditions due to non-linearity in response at high laser powers seen in Supplemental Figure 5, which may lead the maximum photon counts to be an underestimate. However, FRET modeling conducted at <30% donors excited per pulse does not violate any model assumputions nor is it mathematically expected to yield fundamentally different results than at 30% donors excited per pulse. Moreover, simulations run at 10% donors excited per pulse to confirm this expectation showed no significant difference in model output (data not shown). A lifetime value was assigned to each excited particle by generating a random number from an exponential probability density function (τ=2.4 ns; an experimentally determined value of our donor fluorophore). This lifetime value determined how long each excited donor remained excited (in the absence of any FRET process).

At each time point, after the new location of all particles were calculated (using a random walk process), each excited donor neighborhood was checked for acceptors independently. The donor’s neighborhood was defined as a disk with radius of 4-10 nm. In the case where there was an acceptor present in the region of a specific excited donor, the energy of the donor was transferred to the acceptor, according to the rate determined by all donor-acceptor pairs present within the above radial distance. The FRET efficiency for a donor and a single acceptor was calculated from the following equation:

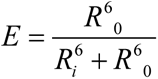

where *R*_0_ is the Förster distance, and *R*_i_ is the distance between donor and acceptor. The FRET efficiency calculation is more complicated when a donor transfers its energy to more than a single acceptor (see “FRET efficiency” below). In both cases, the donor has decayed substantially. For simplicity, the acceptors that received transferred energy from a donor do not get excluded from receiving energy from another nearby donor. In our models, acceptors in this condition are rare and contribute negligibly to the model outcomes. Note that two different phenomena would result in the decay of excited donors: FRET (as explained in this paragraph) and emission (where the excited donor decays to ground state according to its characteristic fluorescence lifetime). The total probability of excited state decay is the sum of probabilities from both processes.

#### FRET Efficiency Calculation (Model)

With the assumption that the concentration of excited donors is lower than the acceptor concentration, we can consider only one donor molecule. In addition, if the orientation factor for dipolar coupling between donor and acceptor is identical for all donor-acceptor pairs, the FRET efficiency equation is as follows, (the pairs are considered to be rotating freely [81]):

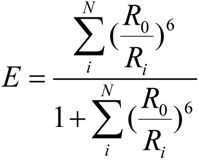

The Förster distance (*R*_0_) of the pair of donor and acceptor fluorophores used for glucan stimulation experiments also matches that used for donors and acceptors in our FLIM FRET experiments, namely *R*_0_= 5.24 nm [82]. The FRET efficiency in this simulation calculates the combination of FRET resulting from acceptors surrounding one excited donor, which are located with the specified radial distance.

### Software

For the RICS and N&B Data presentation and analysis we used the SimFCS Program (www.lfd.uci.edu). The calcium imaging was analyzed using ImageJ. The FLIM-FRET results were analyzed using Symphotime64 software. The BioLayer interferometry analysis was done using BLItz Pro software. Statistical analysis was performed with GraphPad Prism versions 8.2 (GraphPad Software Inc.). DSTORM analysis and FLIM-FRET modeling was performed using MATLAB using our own algorithms (https://github.com/NeumannLab/FRET-Simulation).

## Acknowledgements

The authors declare that they have no conflicts of interest relevant to this work. This research was supported by the University of New Mexico Center for Spatiotemporal Modeling of Cell Signaling (STMC; NIH P50GM085273, AKN) and R01AI116894 (AKN), EUA was supported by fellowships from the STMC and an NIH T32 training grant (NIH T32 AI007538) during the course of this work. We acknowledge the competent technical assistance of Ms. Zinia Pervin in relation to BLI determinations of Dectin-1A/glucan affinity. Glucans used in this study were a generous gift of Immuno Research Inc, which played no role in experimental design, data interpretation or decision to publish. We acknowledge use of the University of New Mexico Comprehensive Cancer Center fluorescence microscopy shared facility, as well as the NIH P30CA118100 support for these cores. We would like to thank the UNM Center for Advanced Research Computing, supported in part by the National Science Foundation, for providing the high performance computing resources used for FLIM-FRET modeling in this work.

## Supplemental Materials

**Supplemental Figure 1.**
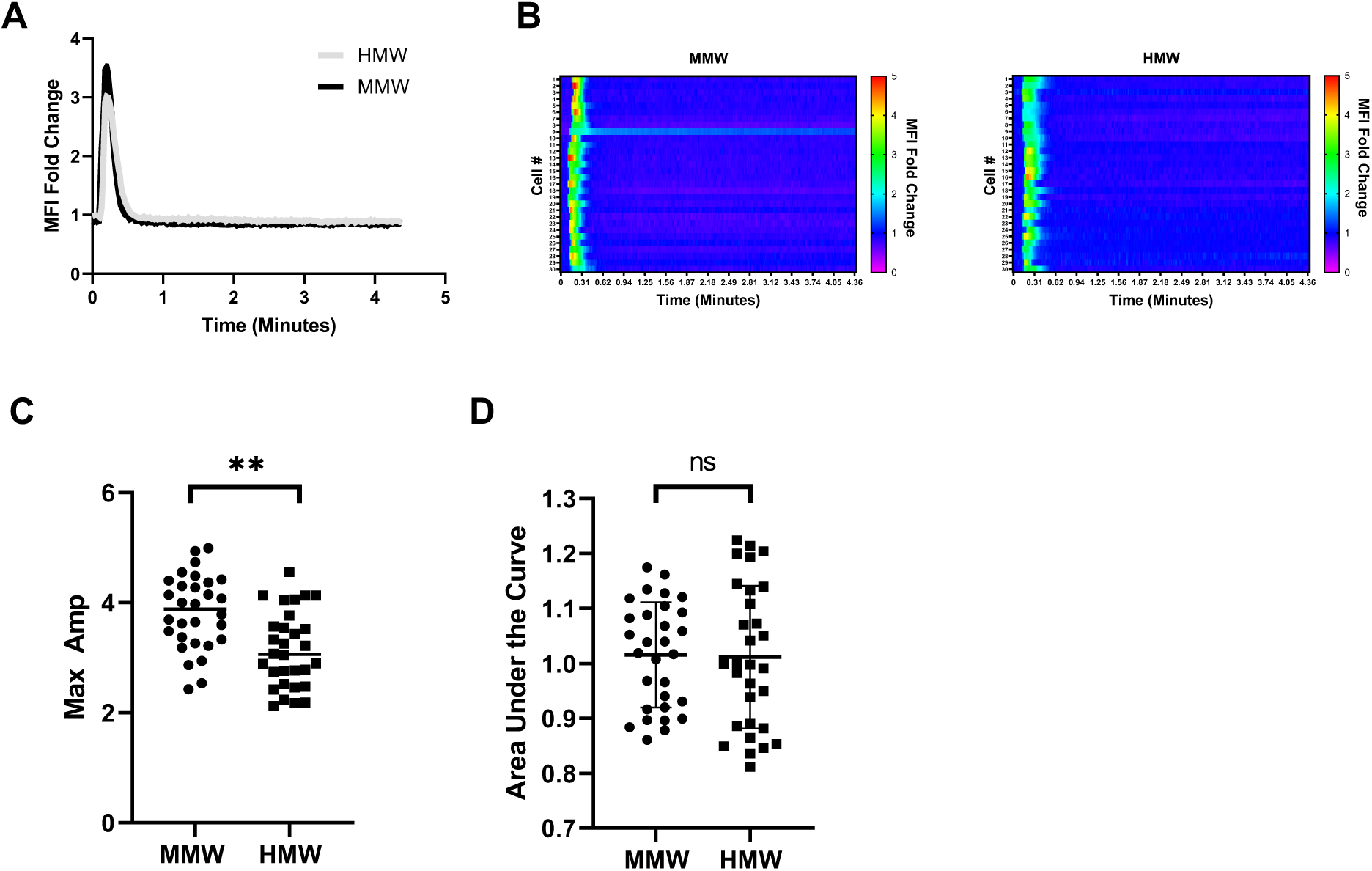
Equimolar glucan stimulation of Dectin-1A exhibits similar response to MMW and HMW glucans. (A) HEK-293 cells stably transfected with Dectin-1A were loaded with Fluo-4 and Cell Tracker Deep Red at equimolar concentrations. Cell Tracker Deep Red was simultaneously loaded in order to normalize for changes in cytosolic volume caused from cell contraction. The mean fluorescence intensity of 30 cells was averaged for Dectin-1A transfected HEK-293 cells stimulated with MMW or HMW glucan at 6.67 nM. (n = 30 per glucan from 3 independent experiments per glucan). Data shown as mean fold change in volume-normalized [Ca^2+^]_intracellular_. (B) Heat maps show relative changes in intracellular calcium concentration of Dectin-1A transfected individual cells upon either addition of MMW or HMW glucan. Each row represents the normalized ratio of Fluo-4 and Cell Tracker Deep Red for a single cell over time. (C) Maximum amplitude of single cells treated with MMW or HMW glucan, as depicted in panel B. Data shown as mean ± SD (n = 30 cells); Welch’s t-test, ** p<0.001 (D) Area under the curve of individual traces of MMW or HMW glucan stimulated cells, as depicted in panel B. Data shown as mean ± SD (n = 30 cells).

**Supplemental Figure 2.**
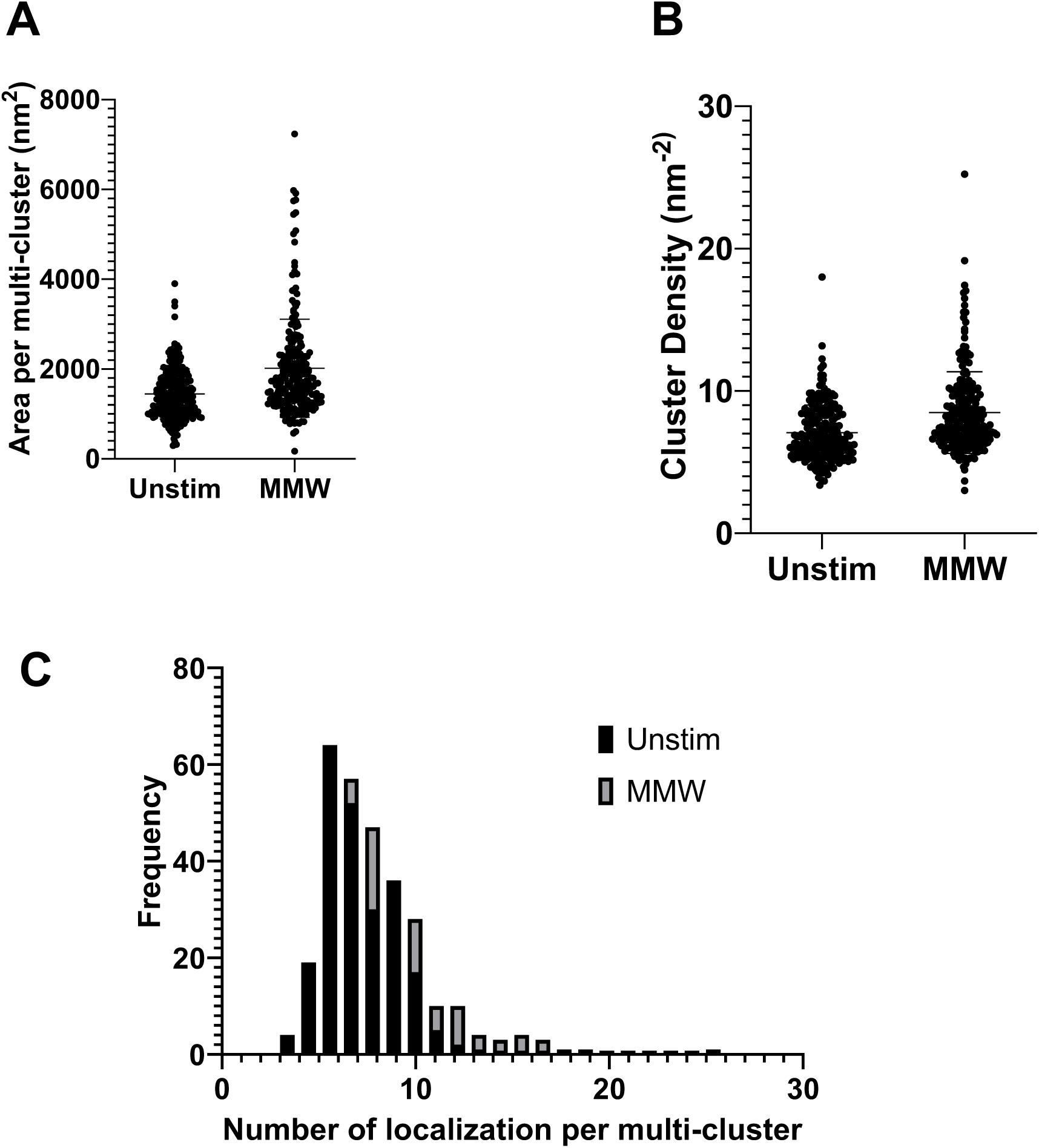
dSTORM clustering analysis shows similar multicluster characteristics in unstimulated and stimulated cells. (A) Areas per multi-cluster in dSTORM analysis of HEK-293 cells expressing Dectin-1A unstimulated or stimulated with MMW. (B) Cluster density of dSTORM analysis of HEK-293 cells expressing Dectin-1A unstimulated or stimulated with MMW. (C) Histogram analysis of the number of localizations of HEK-293 cells expressing Dectin-1A unstimulated or stimulated with MMW. Data shown as mean ± SD (n = 34).

**Supplemental Figure 3.**
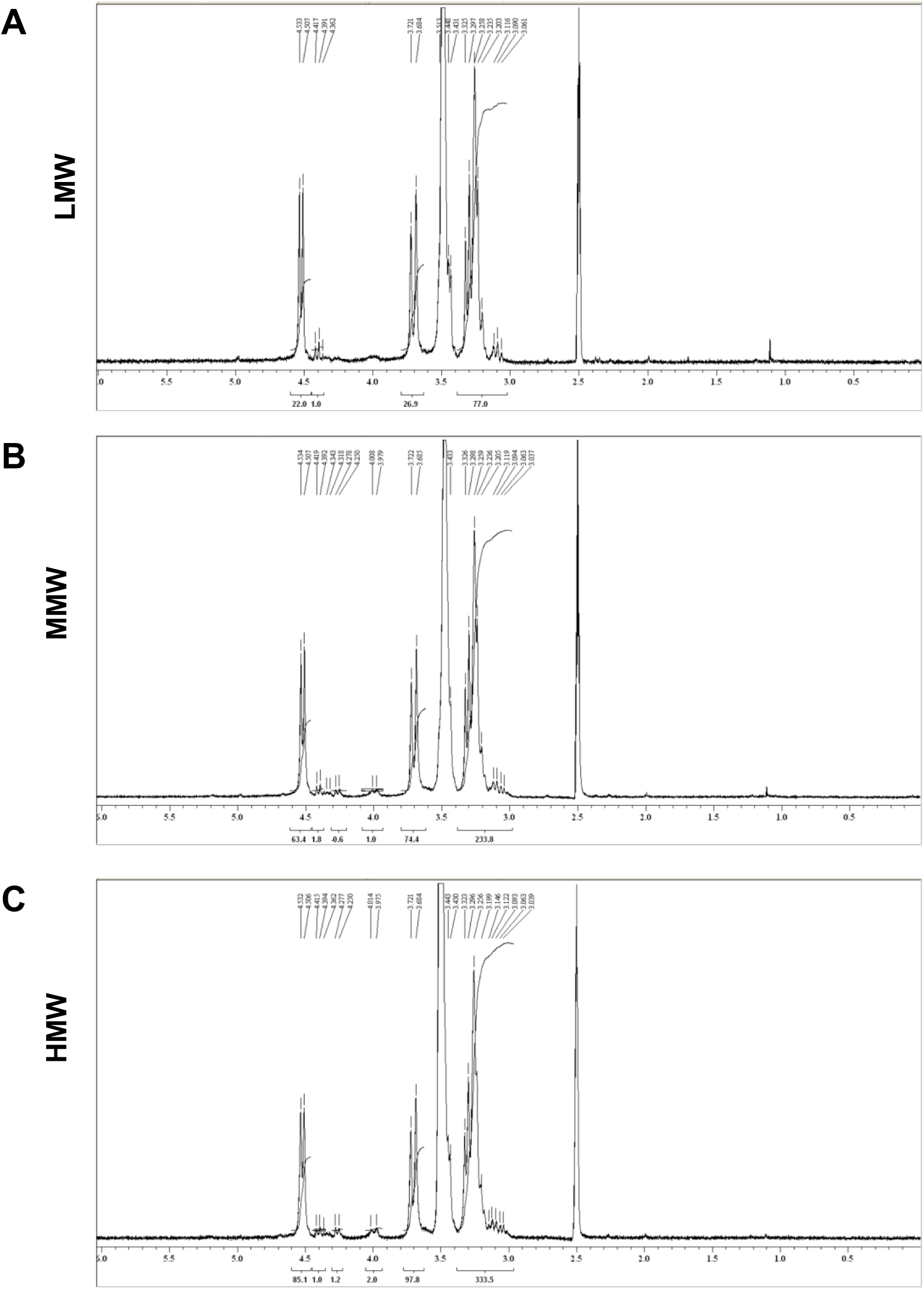
^1^HNMR spectra of glucans used in this study. (A) ^1^H NMR spectrum of LMW, (B) MMW, and (C) HMW glucan.

**Supplemental Figure 4.**
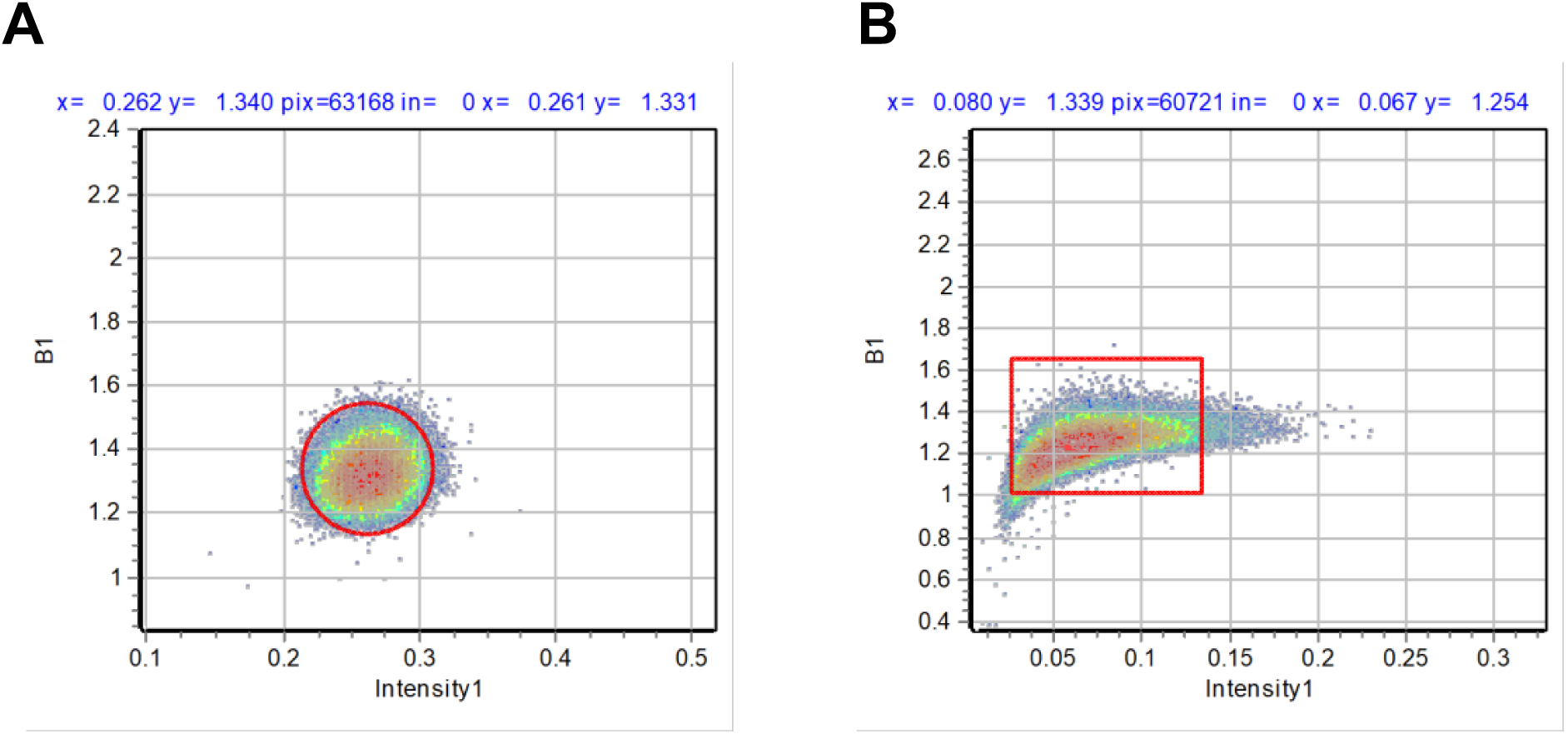
Numbers and Brightness analysis calibration. (A) Brightness vs intensity 2D histogram of purified EGFP with the selected pixels that contribute to the monomers (red circle) in the image. A brightness value of 1.34 was obtained. (B) Brightness vs intensity 2D histogram of purified mEmerald-Dectin-1A with the selected pixels that contribute to the monomers (red box) in the image. A brightness value of 1.34 was obtained.

**Supplemental Figure 5.**
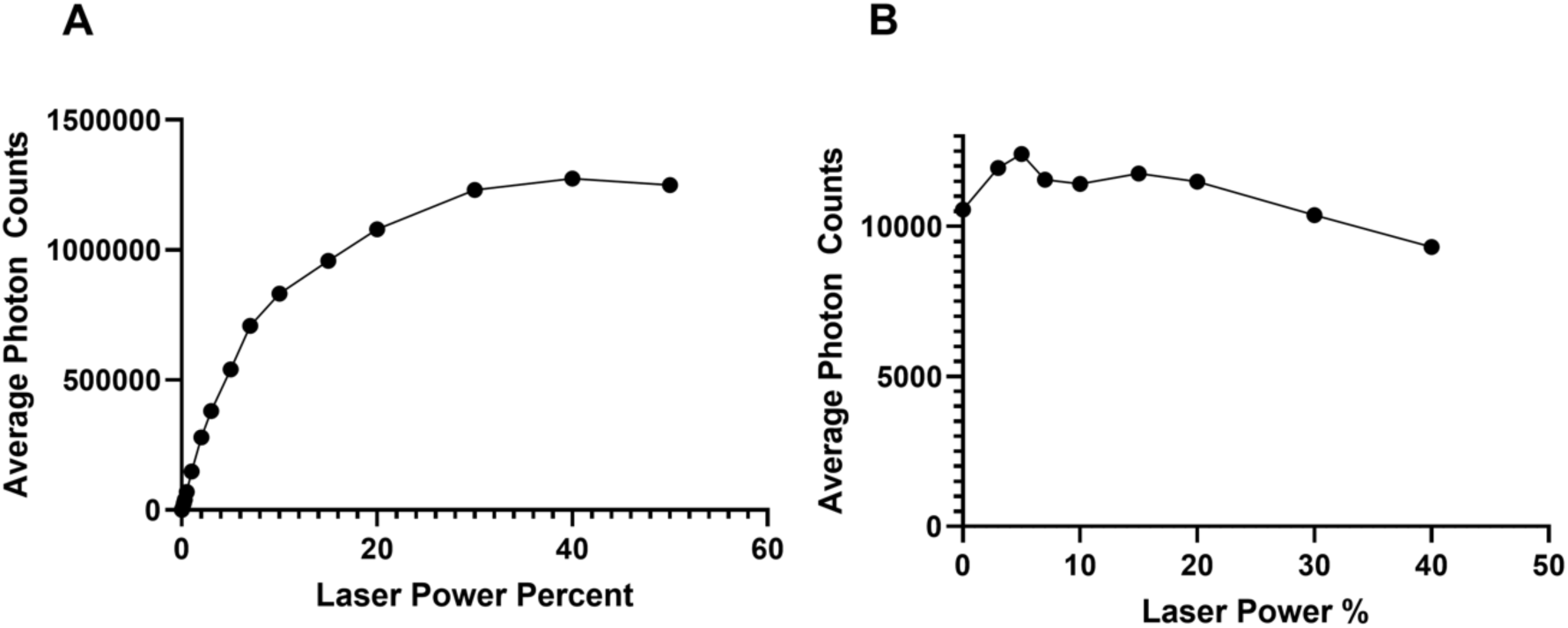
Estimation of fractional donor population excited under FLIM FRET experimental conditions. (A) Average photon counts from cells expressing mEmerald-Dectin-1A were obtained at different excitation laser powers using all other experimental conditions as described in Methods (“Fluorescence Lifetime Imaging Microscopy” section). Maximum photon counts were considered to approximate 100% donor excitation and the percent of donors excited at 3% laser power (the excitation intensity used for experiments) was obtained using the following equation:

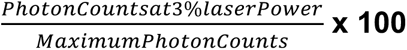

and was used to the parameterize the FRET model. (B) To check for possible photobleaching during acquisition of data in (A), average photon counts were separately acquired (excitation at 0.1% laser power) after each laser power data point in (A) was acquired, from each individual cell. Photobleaching was negligible across the experiments conducted to obtain data in (A).

**Supplemental Table 1.**

| Table of Values |  |  |  |  |  |  |  |  |
| --- | --- | --- | --- | --- | --- | --- | --- | --- |
|  |  | LMW |  | MMW |  | HMW |  |  |
| Max Calcium |  | 1.3 ± 0.1295 |  | 3.4 ± 0.3378 |  | 1.9 ± 0.3837 |  |  |
| Amplitude |  |  |  |  |  |  |  |  |
|  | LMW |  | MMW |  | HMW |  | MMW | MMW |
|  |  |  |  |  |  |  | Denatured | Renatured |
| Kd | 1.93 ± 1.09 |  | 1.62 ± 0.55 |  | 0.43 ± 0.17 |  | 1.02 ± 0.37 | 1.22 ± 1.00 |
| (nM) |  |  |  |  |  |  |  |  |
|  | LMW |  | MMW |  | HMW |  | MMW Denatured |  |
| Min | FRET | Donor- | FRET | Donor- | FRET | Donor- | FRET | Donor- |
|  | Efficiency | Acceptor | Efficiency | Acceptor | Efficiency | Acceptor | Efficiency | Acceptor |
|  | % | % | % | % | % | % | % | % |
| 0 | 83.3 ± | 17.7 ± | 76.5 ± | 15.3 ± | 86.1 ± | 14.5 ± | 87.5 ± | 15.9 ± |
|  | 9.50 | 4.97 | 15.69 | 7.02 | 7.48 | 4.66 | 9.69 | 5.97 |
| 1 | 79.1 ± | 17.1 ± | 77.0 ± | 22.8 ± | 80.1 ± | 19.5 ± | 86.7 ± | 15.8 ± |
|  | 6.31 | 6.48 | 9.05 | 6.54 | 9.05 | 5.49 | 8.36 | 6.06 |
| 2 | 79.1 ± | 17.9 ± | 74.3 ± | 22.1 ± | 78.9 ± | 19.8 ± | 86.2 ± | 15.1 ± |
|  | 8.28 | 5.98 | 9.39 | 8.25 | 8.89 | 6.20 | 8.36 | 6.63 |
| 3 | 78.5 ± | 18.1 ± | 77.1 ± | 23.7 ± | 78.9 ± | 21.7 ± | 86.3 ± | 15.2 ± |
|  | 7.03 | 5.41 | 7.43 | 9.13 | 8.90 | 6.91 | 6.66 | 5.68 |
| 4 | 78.2 ± | 19.5 ± | 72.1 ± | 25.7 ± | 79.5 ± | 22.5 ± | 84.1 ± | 16.1 ± |
|  | 5.85 | 6.89 | 9.05 | 8.87 | 8.12 | 5.40 | 10.29 | 5.51 |
| 5 | 77.9 ±<br>3.96 | 18.8 ±<br>6.16 | 71.9 ±<br>7.76 | 28.2 ±<br>7.12 | 79.5 ±<br>9.14 | 23.7 ±<br>8.16 | 84.5 ±<br>11.45 | 14.1 ±<br>6.70 |
|  | N&B |  |  |  |  | RICS |  |  |
|  | Monomers % | Dimers % | Oligomers % |  | Diffusion Coefficient (μm<sup>2</sup>/s) |  |  |  |
| Unstimulated | 74.0 ± 19.47 | 21.4 ± 18.09 | 4.6 ± 2.39 |  | 1.12 ± 0.50 |  |  |  |
| LMW | 69.5 ± 15.59 | 20.6 ± 9.94 | 9.9 ± 7.15 |  | 0.90 ± 0.46 |  |  |  |
| MMW | 27.5 ± 21.56 | 52.5 ± 15.93 | 20.0 ± 19.49 |  | 0.37 ± 0.24 |  |  |  |
| HMW | 37.5 ± 18.44 | 48.7 ± 14.24 | 13.8 ± 14.35 |  | 0.37 ± 0.18 |  |  |  |
|  | Singlet Cluster<br>Density (nm<sup>-2</sup>) | Multiple Cluster<br>Density (nm<sup>-2</sup>) | Density of<br>Localizations (nm<sup>-2</sup>) |  | Area per<br>cluster(nm<sup>2</sup>) |  |  |  |
| Unstimulated | 0.74 ± 0.46 | 0.59 ± 0.05 | 7.06 ± 1.95 |  | 1444 ± 559 |  |  |  |
| MMW | 0.77 ± 0.46 | 0.66 ± 0.04 | 8.47 ± 2.87 |  | 2016 ± 1093 |  |  |  |
|  |  | FRET Efficiency % |  |  | Donor- Acceptor % |  |  |  |
| Cell Membrane |  | 81.8 ± 14.53 |  |  | 15.7 ± 8.97 |  |  |  |
| SC5314 |  | 84.8 ± 8.67 |  |  | 15.9 ± 6.78 |  |  |  |
| TRL035 |  | 78.8 ± 13.98 |  |  | 25.1 ± 13.06 |  |  |  |
| Glucan Particle |  | 78.8 ± 7.33 |  |  | 29.6 ± 13.36 |  |  |  |

